# Distinct Effects of Aging and Klotho Deletion on the Choroid Plexus

**DOI:** 10.1101/2025.11.07.686549

**Authors:** Zahra Fanaei-Kahrani, Tushar Patel, Christina Valkova, Alexander Gloria, Justine Wagner, Heike Heuer, Markus Schwaninger, Steve Hofmann, Reinhard Bauer, Christoph Kaether

## Abstract

Klotho (Kl) is an anti-aging protein primarily produced in the kidney and the choroid plexus (CP, where it regulates cerebrospinal fluid composition and exerts neuroprotective effects. Here, we investigated the age-dependent consequences of a CP-specific Kl deletion on CP structure and function using mice lacking KL exclusively in CP epithelial cells (Kl^ΔCP^). In control mice, aging markedly disrupted CP architecture and cilia organization both in the lateral (LV-CP) and fourth ventricle (FV-CP). While CP-specific Kl deletion alone caused no major structural changes it induced region- and age-dependent calcification: FV-CP calcification increased in both aged and young Kl^ΔCP^ mice, whereas LV-CP calcification emerged only in older Kl^ΔCP^ mice. Proteomic analysis of the CP and hippocampus revealed mild molecular alterations, suggesting compensatory mechanisms that preserve structural and functional stability despite calcium dysregulation. Consistently, Kl^ΔCP^ mice exhibited no significant behavioral or cognitive deficits. Overall, Kl deficiency sensitizes the CP to age-related calcification prior to overt structural decline, revealing a region-specific and functional link between Klotho, calcium imbalance, and brain aging.

## Introduction

The discovery of the *Klotho* gene (Kl) in 1997 by Makoto Kuro-o and his team opened a new chapter in aging research [1]. Kl quickly gained attention for its remarkable anti-aging properties and its important role in human physiology [2, 3]. Mice deficient in Kl exhibit striking symptoms of accelerated aging, including growth retardation, cognitive impairment, and a drastically reduced lifespan of under 100 days [1]. Studies using Kl-deficient mice have provided important insights into the protein’s neurological functions. These mice display Parkinsonian-like gait abnormalities, reduced Purkinje cell numbers in the cerebellum, memory deficits, and neurodegeneration, highlighting the critical role of Kl in the brain [4]. Interestingly, although the kidney is the primary source of Kl in the body, the protein is crucial for brain functions such as memory and cognition [5, 6]. The primary site of Kl production in the brain is the choroid plexus (CP), which raises a key question: what is the specific role of CP-derived Kl in brain function? Specifically, how does CP-derived Kl affect both the CP itself and other brain regions?

Although the CP accounts for only 0.25% of brain volume [7], it is a critical player in brain protection and homeostasis, CSF production, regulation of circadian rhythms, cognition, neuroprotection, neurogenesis, and inflammatory signaling [8]. Structurally, the CP is a highly vascularized secretory tissue found in all brain ventricles, namely the lateral (LV-CP), third (TV-CP), and fourth (FV-CP). It exhibits morphological diversity, appearing sheet-like in the lateral ventricle and more branched in the third and fourth ventricles. Moreover, the protein composition of the CP differs across ventricles [9]. The CP has an epithelial-like organization, consisting of a monolayer of polarized epithelial cells supported by connective stroma and an underlying fenestrated vasculature that enables exchange between blood and cerebrospinal fluid. These epithelial cells are rich in mitochondria, bear cilia and microvilli, and are interconnected by tight junctions [10]. They also express a wide range of transporters, ion channels, and pumps that differ between the apical and basal membranes, making the CP a metabolically active, highly specialized, and essential regulator of brain physiology [10].

The effects of aging on the CP have been studied, but a comprehensive analysis of how aging influences the LV-CP and FV-CP separately is still lacking. Moreover, it is known that Kl is involved in Ca²⁺ homeostasis in the CP, and virally induced CP-specific Kl deletion has been shown not to alter overall CP structure drastically but to increase inflammation and macrophage infiltration, suggesting a link to immune regulation [11, 12]. Kl expression in the CP is reduced in aged mice [12, 13]. Although some functions of Kl in the CP are known, its precise role in the CP, particularly in the context of aging, remains poorly understood. In this study, we aimed to (i) characterize the structural and functional changes in the CP during aging, with a focus on the LV-CP and FV-CP, and (ii) investigate the effects of CP-specific Kl deletion in comparison to aging. Using behavioral tests, histological and immunofluorescent analyses, and proteomic profiling, we compared young and aged wild-type and CP-specific Kl-deficient (Kl^ΔCP^) mice. Aging disrupted CP structure, including cilia loss—most prominently in the LV-CP—whereas Kl deletion did not reproduce these alterations but modestly accelerated age-associated pathways. Calcification increased with age in the FV-CP and was further enhanced by Kl deletion in young mice, while in the LV-CP it was evident only in aged Kl-deficient mice. Despite increased calcification, CP architecture remained largely intact, suggesting that systemic Kl from the kidney may compensate for local loss. Overall, our findings reveal distinct aging effects on CP structure and a context-dependent role of Kl in regulating CP aging and calcium homeostasis.

## Results

### Generation of mice with a CP-specific Kl-deletion

To generate mice with a CP-specific deletion of *Kl* (Kl^ΔCP^), we crossed floxed *Kl* mice (Kl^fl/fl^, with loxP sites flanking exon 2) with transgenic Slco1c1-creERT2 mice, in which tamoxifen (tam)-inducible Cre-ER is expressed under the control of the Slco1c1 locus, driving recombination specifically in capillary endothelial and CP epithelial cells (Slco1c1-creERT2 [14]) In addition, a YFP reporter allele was crossed-in to monitor Cre activity (Figure 1a). We utilized three genotypes: Kl^fl/fl^;Slco1c1-creER^+/+^;YFP^tg/tg^ as control group 1 (Ctrl1); Kl^+/+^;Slco1c1-creER^tg/+^;YFP^tg/tg^ as control group 2 (Ctrl2); and Kl^fl/fl^;Slco1c1-creER^tg/+^;YFP^tg/tg^ as the experimental knockout group (Kl^ΔCP^). Tam-injections were administered on 4-5 week old mice of all groups daily for 5 days, followed by a 1-week break, and then repeated for an additional 3 days. The successful deletion of Kl in the CP of Kl^ΔCP^ mice was confirmed through Western blot analysis, LC/MS proteomics, and immunohistochemical staining (Figure 1b, Supplementary Figure 1).

**Figure 1.**
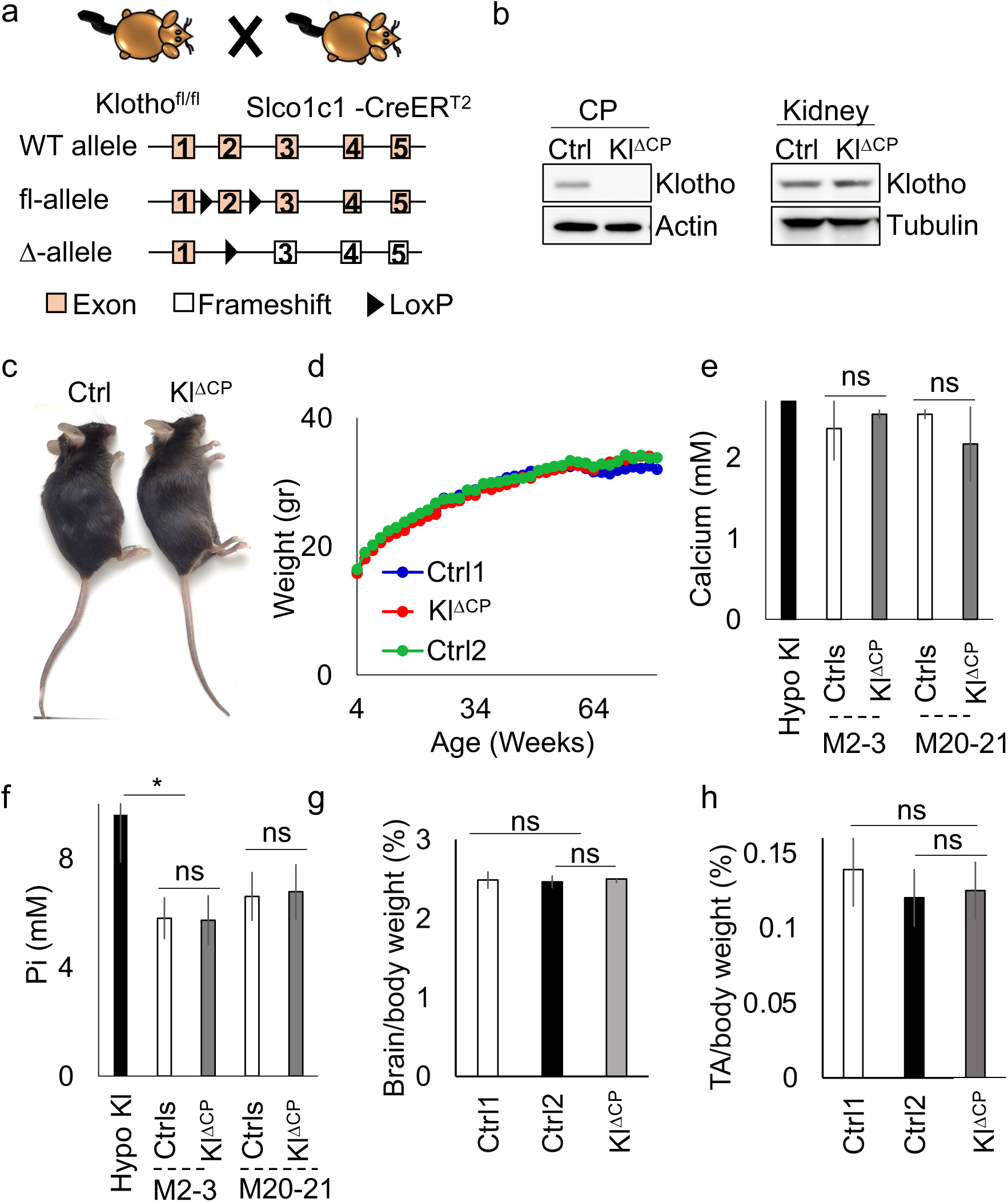
Kl^ΔCP^ mice do not show an apparent phenotype. (a) Breeding scheme for the generation of inducible CP-specific Kl knockout mice. Kl-flox mice were crossed with Slco1c1-creERT2 mice, resulting in Cre-recombinase activation in the CP following TAM injection, and with YFP reporter mice for cre-activity. In Kl^ΔCP^ mice, exon 2 of Kl, flanked by loxP sites, is excised, causing a frameshift and resulting in a nonfunctional, truncated protein. The following genotypes were used: Kl^fl/fl^;Slco1c1-creER^+/+^;YFP^tg/tg^ as control group 1 (Ctrl1); Kl^+/+^;Slco1c1-creER^tg/+^;YFP^tg/tg^ as control group 2 (Ctrl2); and Kl^fl/fl^;Slco1c1-creER^tg/+^;YFP^tg/tg^ as the experimental knockout group (Kl^ΔCP^). (b) Kl^ΔCP^ and Control (ctrl)1 mice were treated with Tam as described in Materials and Methods. At 2-3 months of age, CP and kidney were isolated, lysed, and analyzed by Western blotting using antibodies against Kl and actin or tubulin (c) Representative morphology of Kl^ΔCP^ and control 1 mice at the age of 20-21 months. (d) Body weight measurements over 84 weeks for Kl^ΔCP^ and control mice (n ≥ 9 per genotype). Ca²⁺ (e) and inorganic phosphate (Pi) (f) levels in blood sera of Kl hypomorphic mice, Kl^ΔCP^, and control mice (Ctrls, Ctrl1 and Ctrl2 combined, n ≥ 3 per genotype). (g) Brain/body weight ratio comparison (n ≥ 6 per genotype). (h) TA muscle/body weight ratio at 20–21 months in Kl^ΔCP^ and control groups (n ≥ 9 per genotype). Statistical significance was assessed using a two-sided Student’s t-test; error bars represent ± SD. p > 0.05: not significant (ns), * for (p<0.05).

### Unaltered gross phenotype, survival, and serum biochemistry in Kl^ΔCP^ mice

Kl^ΔCP^ mice were viable and did not exhibit any noticeable behavioral or physical abnormalities. Their survival rates were comparable to those of control littermates, with no increased mortality observed up to 20–21 months of age (Figure 1c, d). Furthermore, blood serum samples were collected from Kl^ΔCP^ and both control 1 and 2 mice at 2–3 months and 20–21 months of age, alongside with samples from 5-week-old Kl hypomorphic mice, known to have abnormalities in Ca^2+^ and phosphate levels [1]. Our results showed a slight increase in serum Ca^2+^ and elevated phosphate concentrations in Kl hypomorphic mice, whereas no significant differences were observed between the Kl^ΔCP^ and control groups (Figure 1e, f). Additionally, the weights of the tibialis anterior (TA) muscle and the brain were measured at 20–21 months and correlated to body weight. No differences in the TA muscle or brain/body weight ratio were detected between the groups (Figure 1g, h), suggesting that the absence of Kl in the CP does not impact these parameters even at advanced age.

### Aging, but not Kl-deletion, affects the morphology and cilia structure of the LV-CP

To assess structural alterations in the CP during aging and after Kl deletion, we performed histological analyses on coronal brain sections. Hematoxylin and eosin (H&E) staining was applied to visualize general tissue architecture and cellular organization. Histological examination of the LV-CP revealed distinct structural differences between young (2–3 months old) and aged (20–21 months old) Ctrl mice. The LV-CP in both young Ctrl and Kl^ΔCP^ mice exhibited well-integrated and uniform architecture, with tightly connected cells forming a cohesive structure. In contrast, aged mice displayed a less organized LV-CP, characterized by areas of cellular separation and detachment (Figure 2a, red arrows). Similar patterns were observed in aged Kl^ΔCP^ mice.

**Figure 2.**
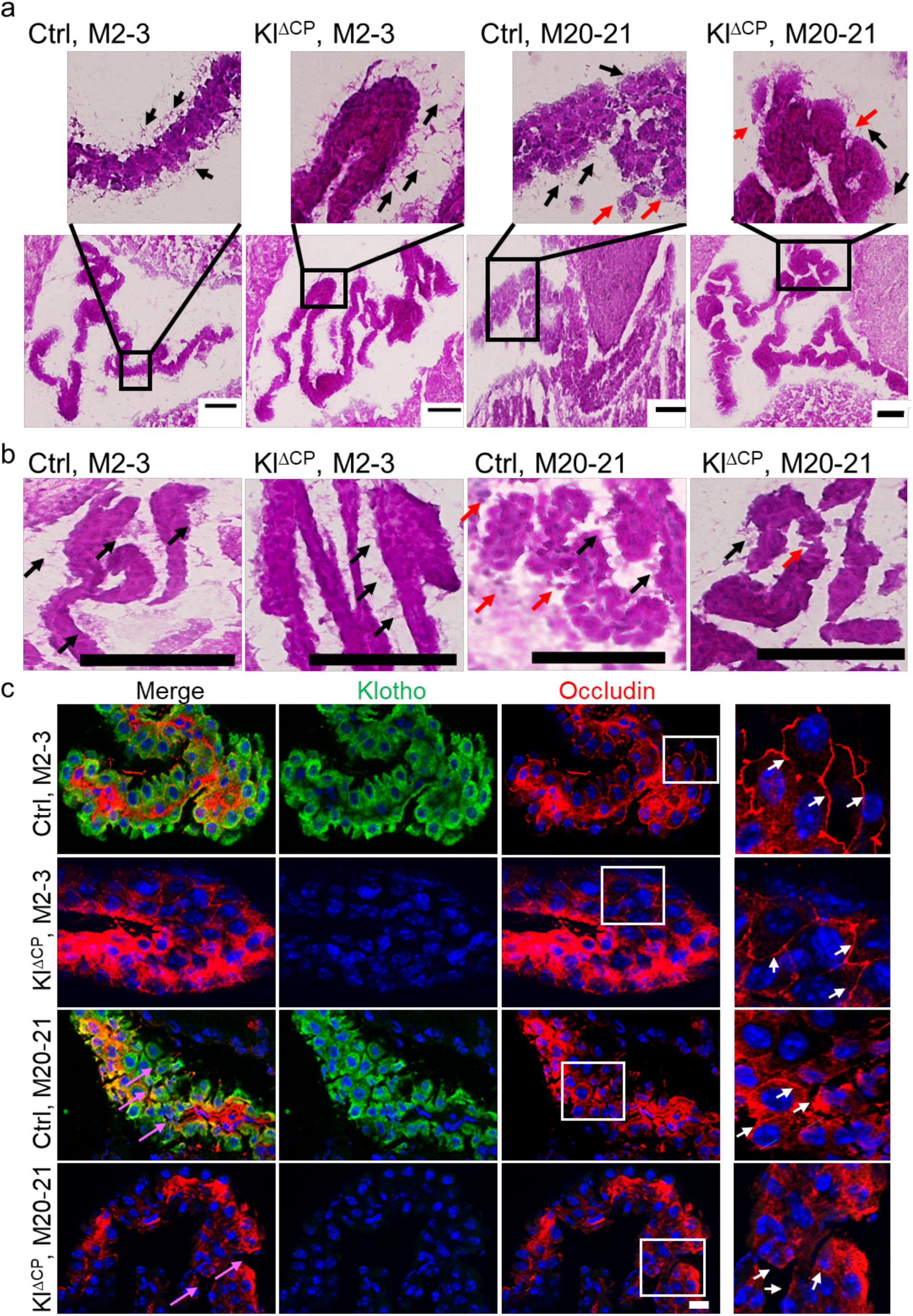
Aging-induced structural deterioration in LV-CP and FV-CP is independent of Kl deletion. Coronal brain sections from control and Kl^ΔCP^ mice at 2–3 months (young) and 20–21 months (aged) were H&E stained and imaged using a Zeiss Axio Scan.Z1 slide scanner microscope, focusing on the LV-CP (a) or FV-CP (b). Ctrl refers to control 1. Black arrows indicate cilia, and red arrows highlight areas of structural deterioration. Scale bar = 100 μm. (c) Immunofluorescent staining of LV-CP for occludin and Kl in control and Kl^ΔCP^ mice at indicated time points. Nuclei are stained with Hoechst (blue). Images were captured using a Zeiss ApoTome microscope. The white arrows indicate the organization of occludin. The pink arrows highlight cell organization and the spaces between cells. The white squares mark enlarged areas in the right panels. Scale bar = 20 μm.

Ciliary morphology further underscored these differences. In young mice, the LV-CP exhibited longer cilia, indicating healthy and functional tissue. [15] (black arrows in Figure 2a). However, aged mice demonstrated a reduction in both ciliary length and density in both aged Ctrl and Kl^ΔCP^ mice (black arrows in Figure 2a). Interestingly, while the absence of Klotho induces premature aging in other tissues [1], it does not show the same effect in the LV-CP of Kl^ΔCP^.

Comparing young and aged FV-CP showed fewer structural changes as seen in the LV-CP. However, signs of structural deterioration (red arrows in Figure 2b) and a reduction in cilia can still be observed with age (black arrows in Figure 2b). These findings suggest that aging exerts a region-specific impact on CP architecture, independent of Kl expression.

Moreover, tight junctions are critical for maintaining the integrity of the blood-cerebrospinal fluid barrier in the CP by tightly sealing adjacent epithelial cells [16]. Occludin, a tight junction marker [16], exhibited a well-organized and continuous pattern at cell-cell borders in both control and Kl^ΔCP^ young mice, consistent with a functional barrier. However, in aged mice, occludin was more irregularly distributed, with gaps and disruptions appearing in their alignment (Figure 2c). This loss of structural cohesion likely underpins the detachment and separation of cells observed in histological analyses (Figure 2b).

### Increased calcification in the CP of Kl^ΔCP^ mice

To investigate the effects of aging and Kl deletion on calcification within the CP, we prepared coronal brain sections and performed von Kossa staining to visualize calcium deposits. We then compared the extent of calcification between Kl^ΔCP^ and control mice, examining both young (2–3 months old) and aged (20–21 months old) animals. The results showed that calcification increased with age, particularly in the FV-CP. Moreover, the deletion of Kl further enhanced the extent of calcification relative to the total tissue area in this tissue (Figure 3a, c). In the LV-CP, this effect was more pronounced at advanced ages, where Kl deficiency led to a marked increase in calcium deposition (Figure 3b, d). These findings are consistent with previous studies suggesting that Kl plays a critical role in calcium and phosphate homeostasis by regulating ion transport and inhibiting ectopic mineralization [17, 18]

**Figure 3.**
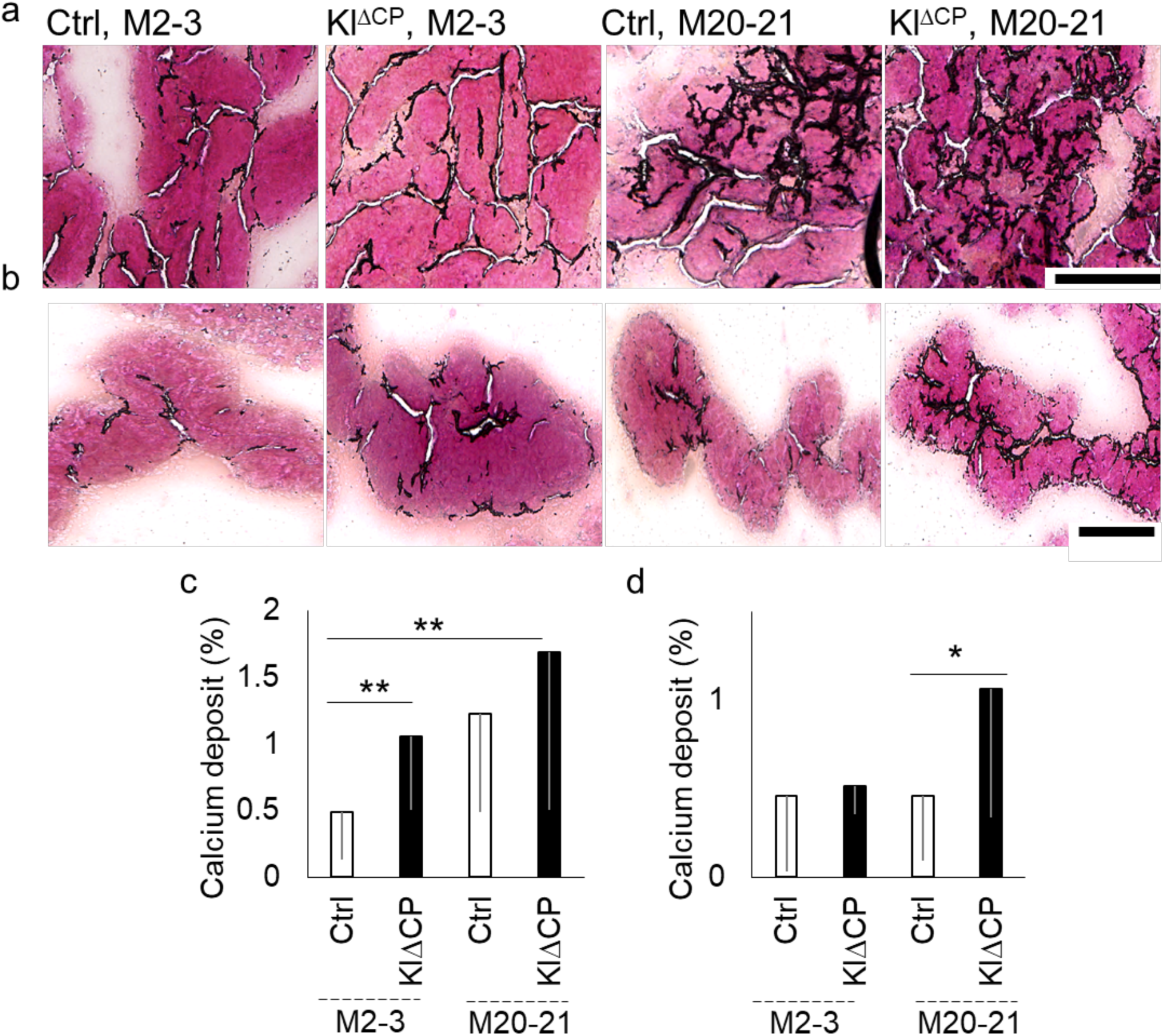
Elevated calcification in CP by specific Kl deletion in the CP and aging. Brain sections from 2-3 and 20–21 months old Kl^ΔCP^ and control mice were subjected to Von Kossa staining and imaged with Zeiss Axio Scan.Z1 slide scanner microscope to visualize calcium deposits in FV-CP (a) and LV-CP (b). (scale bar=100 µm). (c, d) Quantification of calcium deposition in FV-CP (c) and LV-CP (d) normalized to tissue area. (Ctrl: Control, Qty: Quantity). Data was analyzed using a two-sample *t*-test assuming unequal variances. Error bars represent ± SD, Differences were considered not significant (*p* > 0.05, ns). * for (p<0.05), ** for (p<0.01).

### Deletion of Kl in the CP has no effect on the behavior and cognition of young and middle-aged mice

Behavioral experiments, including IntelliCage, rotarod, open field, Morris Water Maze (MWM), hot plate, tail flick, and grip strength, were conducted in two age groups of young (2-5 months) and middle-aged mice (8-12 months) of all three genotypes (Figure 4a). Motor coordination and motor learning were evaluated using the rotarod test, which measures the latency to fall from a rotating rod across repeated trials [19]. All groups demonstrated improvement in performance over successive trials, indicating motor learning. However, no significant differences in latency to drop were observed between Kl^ΔCP^ mice and their control counterparts, suggesting that the deletion of Kl in the CP does not impair motor coordination or learning under these test conditions (Figure 4b, c). Moreover, Kl has been reported to improve learning and memory performance in mice [5]. We therefore next explored the cognitive effects of a deletion of Kl in the CP using an IntelliCage setup [20]. Briefly, the IntelliCage system is an automated behavioral testing setup that allows continuous monitoring of group-housed mice in a specialized polycarbonate cage containing four operant corners. Each mouse is identified by a subcutaneously implanted RFID tag, enabling precise, individual tracking of activity over a 17-day testing period. The system records multiple cognitive and behavioral parameters, including exploration time, movement patterns, spatial learning, relearning performance, and corner visit frequency. Relearning capacity, assessed using a relearn score derived from corner navigation tasks, showed no significant differences between Kl^ΔCP^ and control mice (Figure 4d, e). While Kl^ΔCP^ mice exhibited improved relearn scores compared to one control group (Ctrl2), the difference to Ctrl 1 was not statistically significant. These findings suggest that CP-derived Kl does not significantly contribute to learning or relearning (Figures 4d and e). Next, a water maze test was conducted to assess spatial learning and memory in mice. The test measured their ability to locate a hidden platform over four consecutive days, followed by a probe trial to evaluate memory retention based on the time spent in the target area where the platform was previously located [21]. All groups showed improved performance by reducing the time to find the platform, indicating enhanced spatial learning (Figure 4f, g). However, no significant differences were observed between Kl^ΔCP^ and control mice in learning or memory retention, as shown by similar platform crossings in the probe trial. These results suggest that Kl deletion in the CP does not affect spatial learning or memory. Additional behavioral tests involved open field to assess anxiety level and exploratory behavior (suppl. Figure 2a, b), hot plate (suppl Figure 2c, d) and tail flick response to test nociception (suppl. Figure 2e, f). Finally, grip strength to assess skeletal muscle strength with 2- and 4-paw was determined (suppl. Figure 2g-j). All tests were performed at 2–5 months and 8–12 months. None of the tests showed a significant difference between Kl^ΔCP^ and control mice. Taken together, all tests indicated that the deletion of Kl in the CP does not significantly impact cognition, spatial learning, memory, pain response, general motor coordination, or muscle strength in mice.

**Figure 4.**
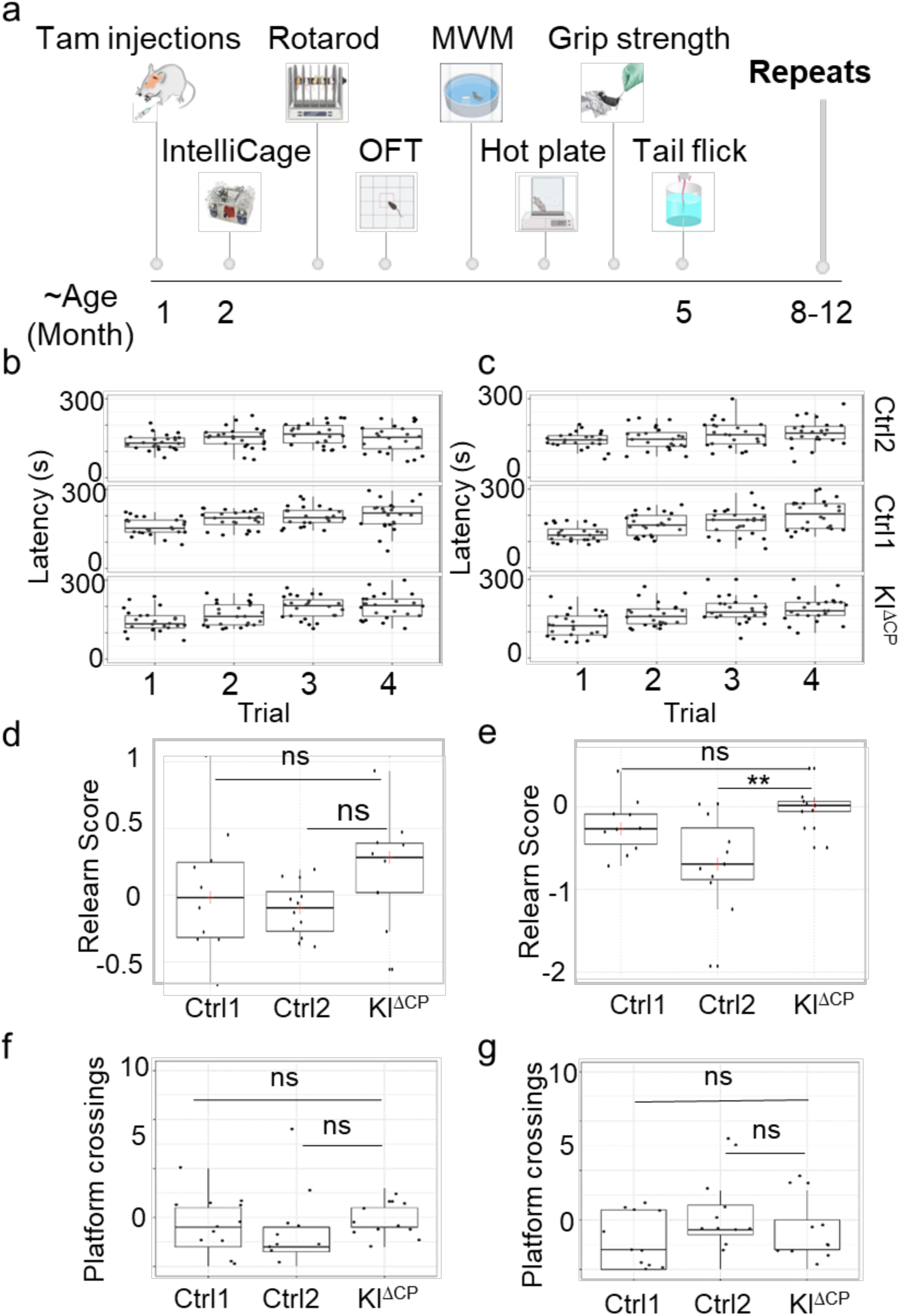
Deletion of Kl in the choroid plexus does not affect behavioral traits such as motor coordination, cognition, or spatial memory in mice. (a) Schematic of behavioral experiments partially created using BioRender.com. Behavioral tests were performed following TAM injections at the age of 2–5 months and repeated at 8–12 months. (b, c) Motor coordination and learning were measured using the accelerating rotarod test in (b) 2–5 months old and (c) 8–12 months old mice. After habituation, the rotarod test was conducted for four consecutive days with an acceleration from 4 to 40 rpm over 5 minutes. The time period before the mouse fell off or passively rotated was recorded. Tests were conducted twice per day with a break of one hour between sessions. (d, e) Relearn scores were calculated as logarithm (ln) (Place Errors/Reversal Errors) to assess relearning capacity during the IntelliCage experiment in (d) 2–5 months and (e) 8–12 months old mice. Higher relearn scores indicate better relearning efficiency. (f, g) The Morris water maze was used to evaluate spatial memory by measuring the number of crossings over the target quadrant (former platform location) during the probe trial in (f) 2–5 months old and (g) 8–12 months old mice, after a 4-day learning phase; All experiments were conducted on n=12 female mice per genotype. Statistical analysis was performed using RM-ANOVA combined with Cohen’s d effect size, and error bars represent standard deviation. Each dot represents data from an individual mouse. A p-value > 0.05 indicates no significant difference: ns; MWM: Moris Water Maze, OFT: Open field, Ctrl: Control.

### Kl deletion in the CP leads to differential protein expression across ages

We next analyzed the proteomic changes in the CP and in the hippocampus (HC) following Kl deletion and assessed their association with age. To this purpose, we conducted separate proteomic analyses on FV-CP and LV-CP, as well as on the HC of control (Ctrl1, Ctrl2) and Kl^ΔCP^ mice at 2-3, 5-6, 14-15, and 20-21 months (Figure 5a). The analysis was performed with 3–5 biological replicates per condition. At each time point, FV-CP, LV-CP, and HC were isolated separately from individual mice under aseptic conditions, ensuring each tissue served as an independent biological replicate for accurate and reliable assessment. Principal Component Analysis (PCA) revealed distinct proteomic profiles for FV-CP, LV-CP and HC, demonstrating reproducibility and specificity of our analysis (Suppl. Figure 3). We confirmed the successful deletion of Kl in Kl^ΔCP^ mice across all ages in LV-CP and FV-CP (Suppl. Figure 1a, b). To explore the age-specific effects of Kl deletion, we performed differential expression analysis for each age group. This was necessary because different life stages, from early development to aging, are associated with unique physiological processes that may influence protein expression. Our analysis excluded proteins that showed differential expression between control sets at any time point. This exclusion was based on DESeq analysis (using the DESeq2 library in R version 1.40.2) with a p-adjusted significance threshold of 0.05. We then compared Ctrl1 and Ctrl2 separately against Kl^ΔCP^ mice, using a log_2_ fold-change cutoff of ±0.6 and a Q-value of <0.05. Results were visualized through volcano plots to highlight upregulated and downregulated proteins in FV-CP and LV-CP in Kl^ΔCP^ versus controls (Figure 5b-c, Suppl. Excel Table S3, 4). The comparison Kl^ΔCP^ /Ctrl2 is shown in Suppl. Figure 4. To identify proteins that are consistently dysregulated across all age groups, we compared the differentially expressed proteins in both Ctrl1 and Ctrl2 relative to Kl^ΔCP^ across multiple age cohorts. This comparison was visualized using a Venn diagram (Figure 5d, e, underlying data in Suppl. Excel Table 3, 4). Although no single protein was consistently dysregulated across all age groups, some proteins were shared between two or more age groups, suggesting that the proteomic response to Kl deletion is complex and highly age dependent.

**Figure 5.**
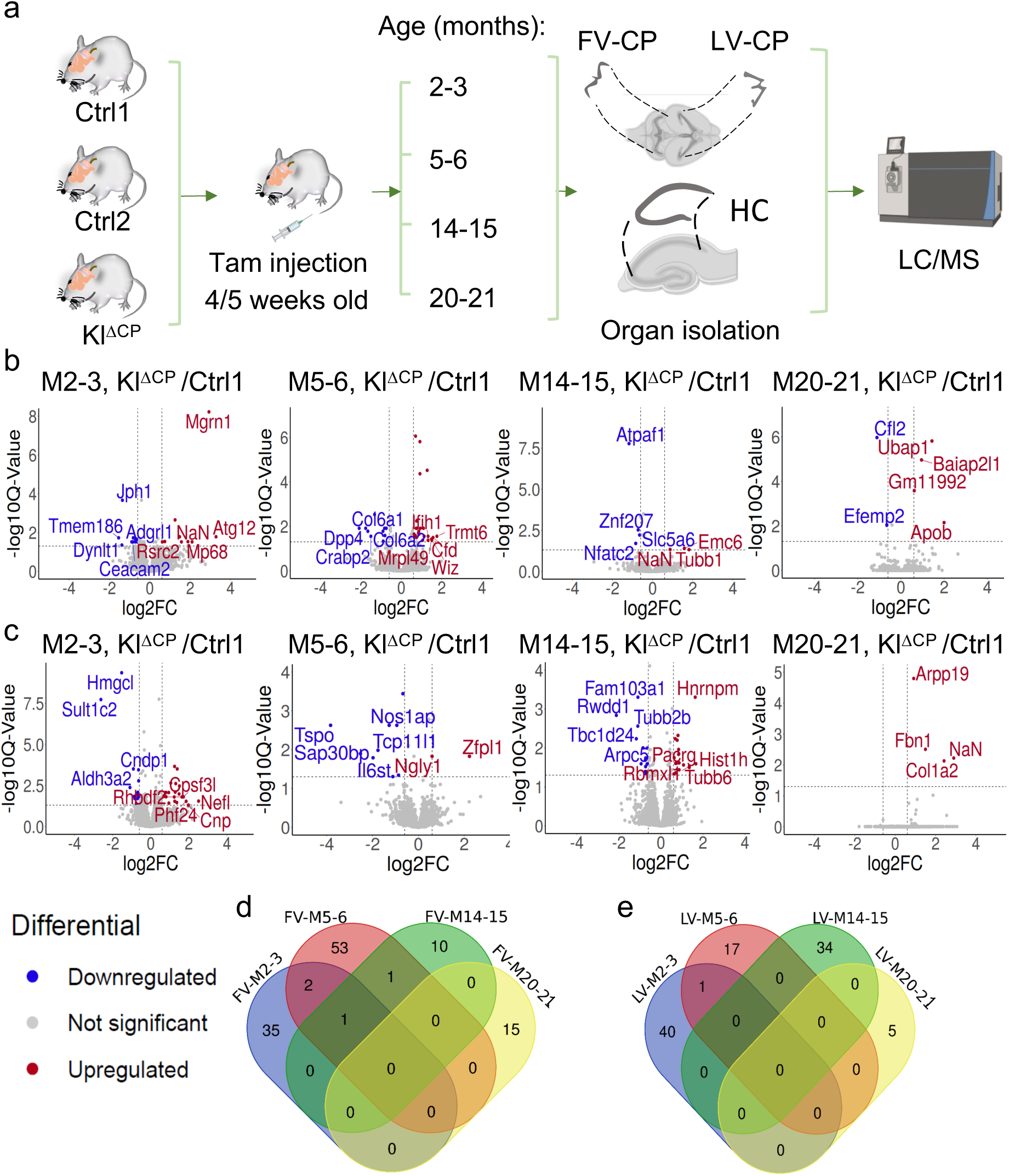
Age-specific effects of Kl deletion in LV-CP and FV-CP. (a) Schematic diagram showing organ isolation and preparation for proteomics analysis. CP and HC from various genotypes and age groups were isolated at specific time points and subjected to LC/MS analysis. The image was partly created with BioRender.com. FV: fourth ventricle, LV: lateral ventricle, HC: hippocampus. (b, c) Volcano plots showing the differential expression of proteins in (b) the FV-CP or (c) the LV-CP of Kl^ΔCP^ mice compared to Control 1 (Ctrl1) at the indicated ages. M, month. Blue dots represent downregulated, red dots upregulated proteins in Kl^ΔCP^ mice, Log_2_FC: Log_2_ Fold Change (for detailed data, refer to the Suppl. Excel Table S1, 2). The top 5 dysregulated proteins according to Q-value and log₂ fold change are shown in the volcano plots. (d-e) Venn diagrams illustrating the overlap of differentially expressed proteins at indicated ages in months (M) in the CP of the (d) FV and (e) LV (for detailed data see Supp. Table 3, 4). To assess the overall impact of Kl deletion in the CP across different ages, we performed a comprehensive analysis by pooling the common proteins that showed significant changes when comparing Kl^ΔCP^ mice with both control groups (Ctrl1 and Ctrl2) individually. By analyzing the proteins that were significantly altered in the Kl^ΔCP^ versus each control group separately, we could identify those consistently affected by Kl deficiency throughout the lifespan. If a protein showed consistent changes in both comparisons, its expression is likely to have been directly impacted by the Kl deletion in the CP. The proteins identified through this collective enrichment analysis are listed in Table 1.

**Table 1.**
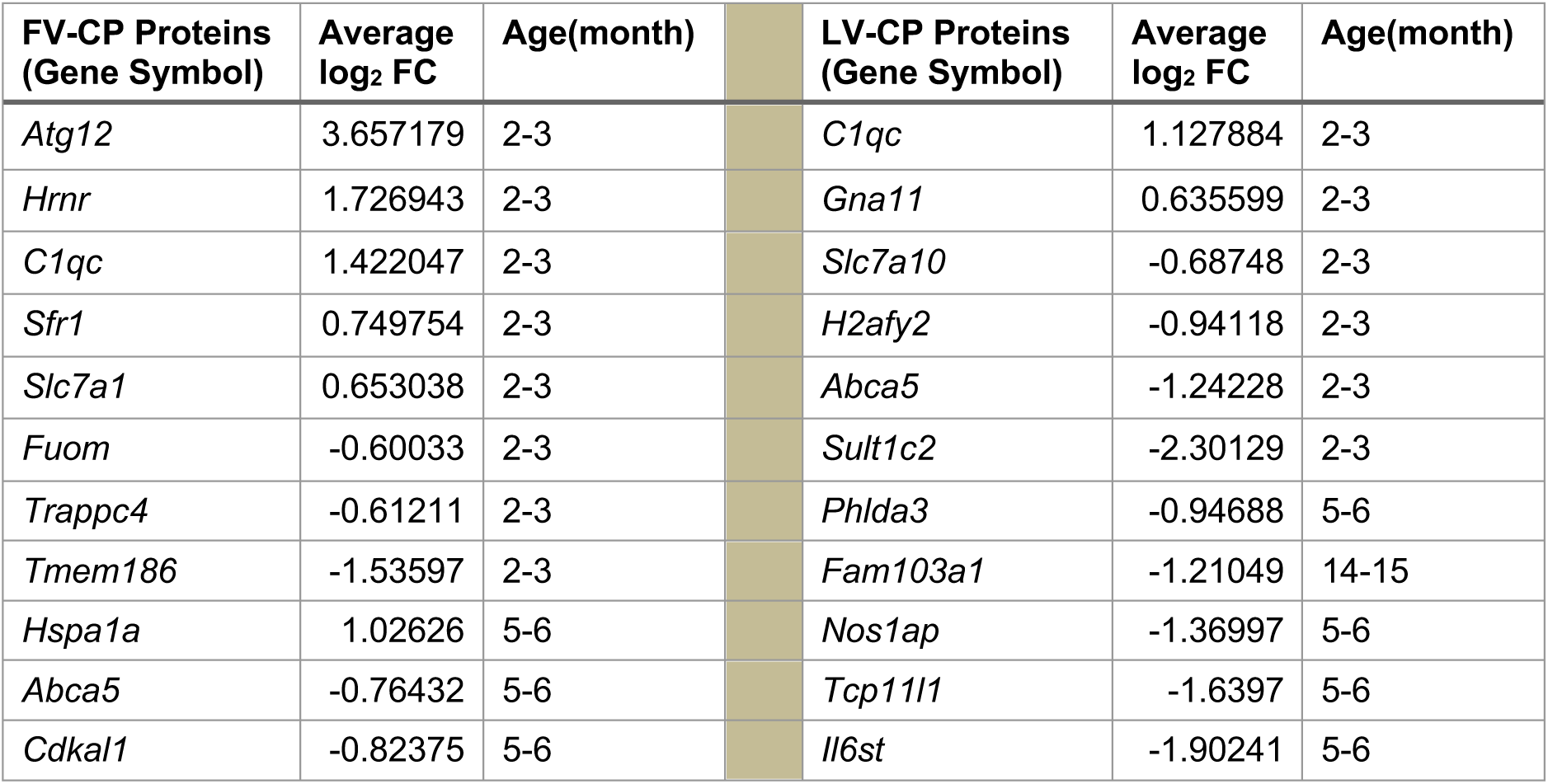
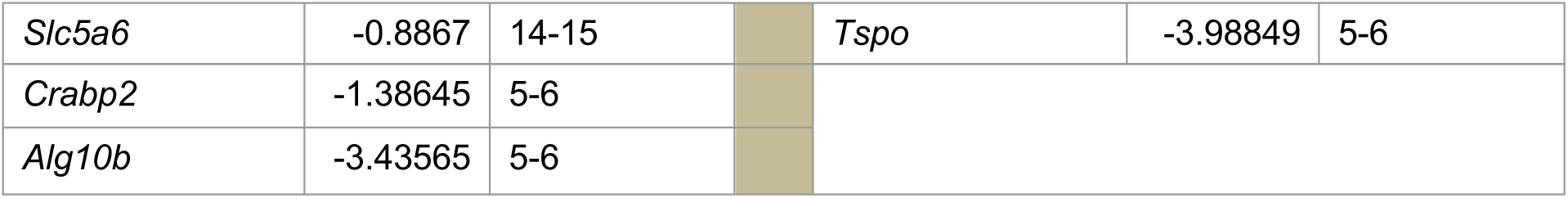
List of common proteins differentially expressed in FV-CP and LV-CP when comparing Kl^ΔCP^ to Ctrl1 and Kl^ΔCP^ to Ctrl2. The average Log_2_ Fold Change (FC) represents the mean of the Log_2_ FC values from comparisons of Kl^ΔCP^ with Ctrl1 and Kl^ΔCP^ with Ctrl2. Positive and negative values indicate that protein levels were higher and lower in Kl^ΔCP^ relative to controls, respectively.

Among the identified proteins upregulated in the FV-CP of Kl^ΔCP^ mice, C1qc is known for its roles in local immune and complement responses [22]. Abca5 and Slc5a6 were downregulated in Kl^ΔCP^ mice and were also detected before in CP transcriptomic and proteomic atlases, suggesting potential roles in lipid and vitamin transport [23, 24]), respectively. The remaining candidates (Atg12, Trappc4, Cdkal1, Crabp2, Alg10b, etc.) are present in CP expression databases (https://www.proteinatlas.org) but currently lack functional characterization in the CP.

In the LV-CP, Il6st and C1qc, down- and upregulated in Kl^ΔCP^ mice, respectively, are functionally involved in CP inflammatory or immune responses [25] [22]. Most other proteins in our dataset (Gna11, Abca5, Nos1ap, Slc7a10, H2afy2, Sult1c2, Phlda3, Fam103a1, Tcp11l1) are detected in CP transcriptome/proteome datasets but published studies defining a CP-specific role are lacking (https://www.proteinatlas.org/). Tspo, typically associated with CP inflammation and calcification [26], was reduced in Kl-deficient CP despite increased calcification. This reduction may reflect a compensatory response aimed at limiting further mineral deposition by modulating mitochondrial activity or the production of reactive oxygen species.

### Age-correlated proteomics analysis of FV-CP and LV-CP: accelerated aging signatures and disrupted homeostasis in Kl^ΔCP^ mice

Next, the proteome data were analyzed to identify proteins whose expression progressively changed with age as a result of Kl deletion in the CP. For this analysis, we first calculated the average log₂ fold change between Kl^ΔCP^ and both control groups (Ctrl1 and Ctrl2) for each protein. Using these averaged values, we then performed a Pearson correlation analysis across four age groups (2–3, 5–6, 14–15, and 20–21 months) to assess how protein expression in Kl^ΔCP^ mice changes with age relative to controls. Correlation coefficients were computed for all proteins using the *cor()* function in R. In this analysis, a positive correlation indicates that protein expression in Kl^ΔCP^ mice increased as the animals aged relative to controls, whereas a negative correlation suggests reduced expression with advancing age. Proteins showing strong correlations (r > 0.9 or r < −0.9) were selected for further analysis (Figure 6a–b).

**Figure 6.**
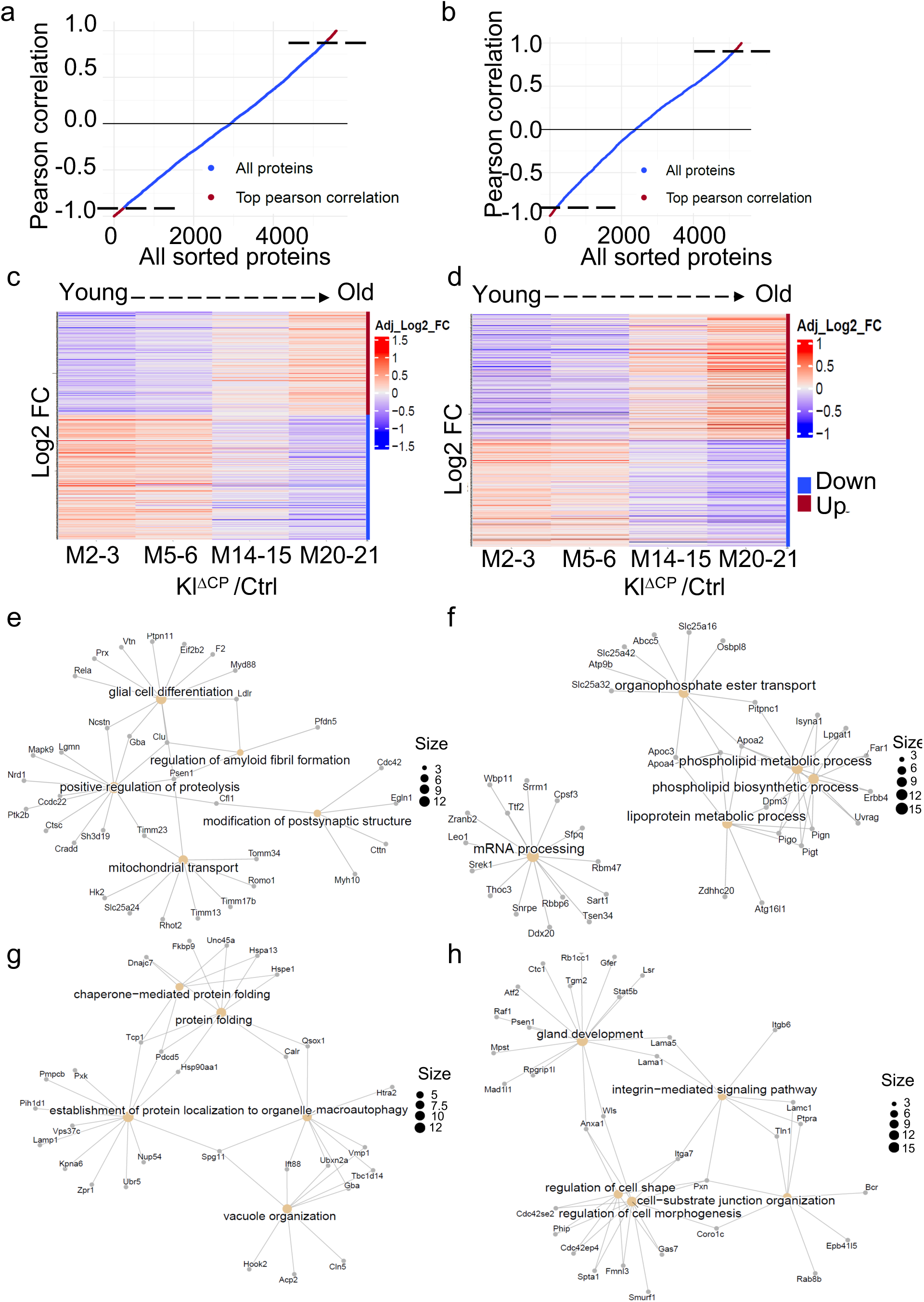
Kl deletion in the CP accelerates aging signatures and activates compensatory or disease-related pathways. (a, b) Pearson correlation analysis of differential protein expression (Kl^ΔCP^ vs. controls) across all timepoints (2-3, 5-6, 14-15, and 20-21 months) in (a) FV-CP and (b) LV-CP. Proteins with Pearson correlation coefficients >0.9/<-0.9 are displayed in red and separated with dotted lines. (c, d) Heatmap of proteins differentially expressed in Kl^ΔCP^ mice versus controls across age groups as indicated. ‘Down’ represents proteins whose expression decreases with age in Kl^ΔCP^ mice, and ‘Up’ indicates increased expression. (c) FV-CP and (d) LV-CP, with log_2_FC representing the average log_2_FC from comparisons of Kl^ΔCP^ /Ctrl1 and Kl^ΔCP^ /Ctrl2. (for detailed data, refer to the suppl Excel table S3, S4) (e, f) Cnet plots showing enriched pathways (enrichGO function from the ClusterProfiler package (version 4.8.3 in R) in FV-CP following Kl deletion, displaying (e) Proteins showing reduced expression with age (negatively correlated) in KlΔCP mice, and (f) proteins showing increased expression with age (positively correlated) in KlΔCP mice (g, h) Cnet plots for enriched pathways in LV-CP, showing (g) downregulated and (h) upregulated candidates with aging in Kl^ΔCP^ mice. Yellow circles represent biological pathways, grey circles represent specific proteins, and grey lines indicate correlations between proteins and pathways. The size of the black dots in the legend reflects the number of processes represented.

These proteins were visualized in heatmap plots for both the LV-CP and FV-CP across all groups (Figure 6c, d), enabling us to identify proteins regulated by Kl during aging. In the FV-CP, 481 out of 5,580 detected proteins showed strong age-related changes in Kl^ΔCP^ mice relative to controls (correlation coefficient > ±0.9). Among these, 218 proteins displayed increased expression with age, whereas 263 proteins showed decreased expression with age. Similarly, in the LV-CP, 402 out of 5,341 proteins showed age-related changes, with 217 increasing and 185 decreasing in expression (Figure 6d, suppl Excel table S3, S4). To explore the biological processes and pathways impacted by CP-Kl deletion during aging, we conducted Gene Ontology (GO) enrichment analysis and visualized them using Cnet plots (Figure. 6e-h). In the FV-CP of Kl^ΔCP^ mice, the proteins whose expression decreased with age were primarily associated with amyloid fibril formation, positive regulation of proteolysis, modification of postsynaptic structures, mitochondrial transport, and glial cell differentiation (Fig. 6E), suggesting that Kl normally supports the homeostasis of these processes. In contrast, proteins that increased in expression with age were linked to lipoprotein and phospholipid metabolism, organophosphate ester transport, and mRNA processing (Fig. 6F), implying that Kl may normally act to restrain or fine-tune these pathways, whose enhancement could contribute to age-related dysfunction. In the LV-CP, proteins that decreased in expression with age in Kl^ΔCP^ mice were associated with protein folding, protein localization to organelles, macroautophagy, and vacuole organization, while those that increased were linked to integrin-mediated signaling, cell-substrate junction organization, and regulation of cell morphology, gland development (Figure 6g–h). The increased expression of integrin and laminin proteins (Itga7, Itgb6, Lamc1, Lama1, Lama5, Tln1, Ptpra) suggests enhanced cell–ECM interactions. Also, proteins involved in cytoskeletal organization and cell–substrate junctions (Pxn, Coro1c, Gas7, Fmn13, Spta1, Cdc42ep4, Cdc42se2, Phip, Anxa1) were also elevated. Based on previous studies, these pathways and proteins have been implicated in extracellular matrix remodeling and mineralization processes [27, 28], suggesting that their increased expression upon KL deletion may contribute to, or be associated with, the observed calcification in LV-CP in older ages. Notably, both the FV-CP and LV-CP showed a decline in protein folding and proteostasis-related pathways, underscoring Kl’s crucial role in preserving protein homeostasis during aging. Nonetheless, region-specific differences highlight that Kl deletion influences aging pathways differently across CP subregions.

### Impact of Choroid Plexus Kl deletion on the hippocampal proteome across age

Next, we performed a proteomic analysis of HC from Ctrl1, Ctrl2, and Kl^ΔCP^ mice at 2-3, 5-6, 14-15, and 20-21 months. First, proteins that were differentially expressed between Ctrl1 and Ctrl2 (cutoff of Q value ≤ 0.05 and log_2_ fold change ≥ +0.6 or ≤ −0.6 were identified and subtracted from the dataset. We then identified proteins that were differentially expressed across the respective ages in Kl^ΔCP^ mice compared to both controls, applying a cutoff of Q value ≤ 0.05 and log_2_ fold change ≥ +0.6 or ≤ −0.6. The results of these comparisons are visualized as heatmap in Fig. 7 and in Supplementary Figure 5 as volcano plots.

**Figure 7.**
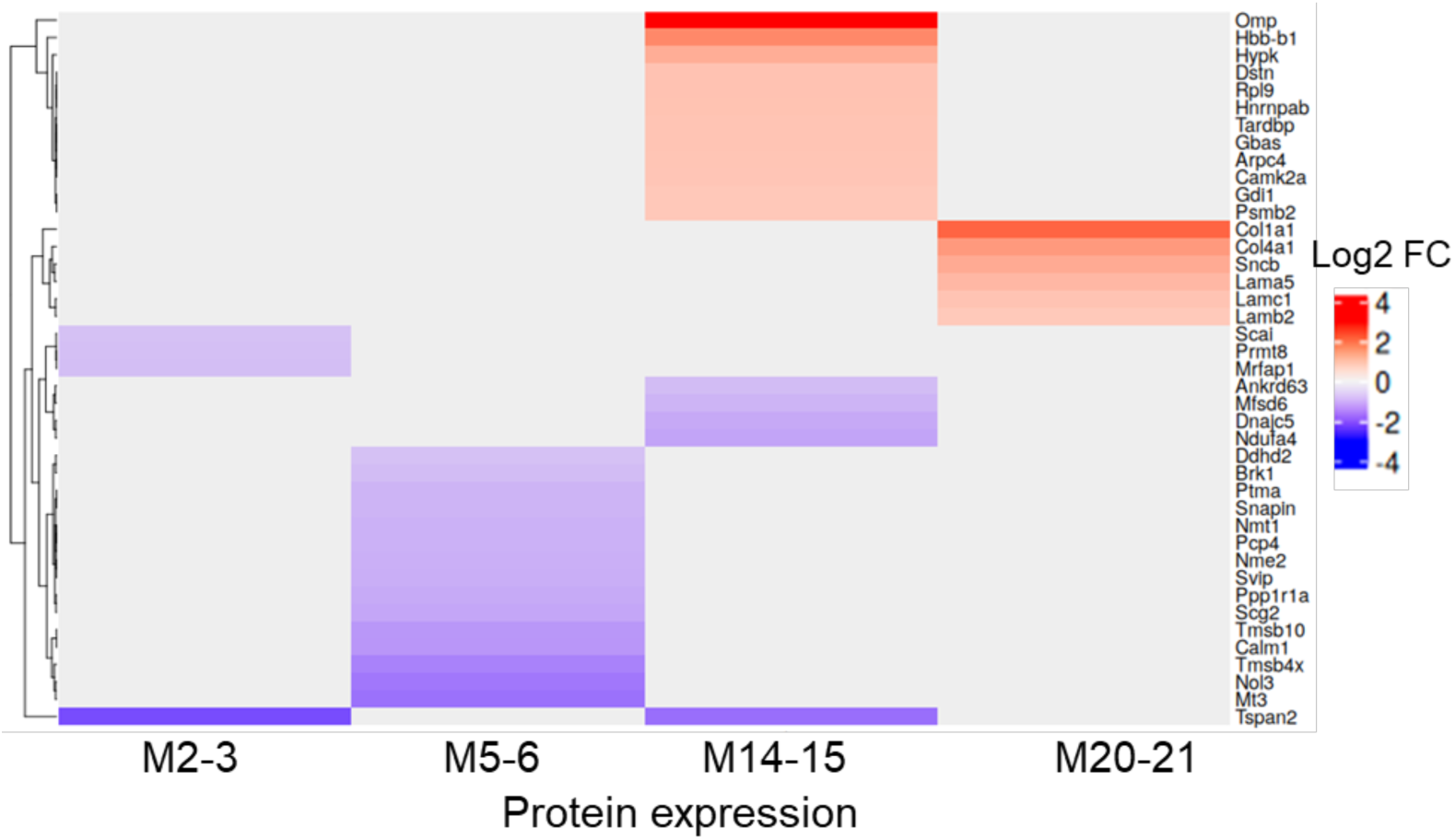
Subtle, age-specific protein expression changes in the HC following Kl deletion in the CP. Heatmap illustrating differentially expressed proteins in the hippocampus of Kl^ΔCP^ mice compared to Ctrl1 and Ctrl2 at ages 2-3, 5-6, 14-15, and 20-21 months (M). Proteins labeled as “upregulated” are upregulated in Kl^ΔCP^ compared to both controls, while “downregulated” indicates downregulation in Kl^ΔCP^ relative to both controls. log_2_FC refers to the mean of the log_2_FC values obtained by individually comparing Kl^ΔCP^ with each control group (Ctrl1 and Ctrl2).

To further explore these patterns, we performed KEGG enrichment analysis on the down and upregulated proteins across all age groups combined, using Enrichr (https://maayanlab.cloud/Enrichr/). This analysis highlighted a notable upregulation of extracellular matrix (ECM)-related proteins, including Col1a1, Lama5, Lamb2, Col4a1, and Lamc1 (Supplementary Table 5, 6, supplementary Figure 5). Also, the absence of Kl in the CP appears to trigger both neurodegenerative and compensatory responses concurrently. Specifically, we observed a decrease in proteins associated with long-term potentiation (LTP), such as Camk2a, Araf, and Calm1, alongside a reduction in proteins linked to neurodegenerative pathways such as Ndufa4. Ndufa4 is known to play a role in AD pathogenesis [29], suggesting a potential protective mechanism by the brain to mitigate the effects of Kl deficiency.

## Discussion

In this study, we investigated the effects of a specific Kl deletion in the CP of mice. Our results show that aging leads to CP disruption and loss of cilia, particularly pronounced in the LV-CP. We also observed an increased calcification, which was more evident in the FV-CP. A similar calcification was observed in aging CP of humans [30]. Interestingly, CP-specific Kl deletion did not affect the aging phenotype, as the overall CP structure and cilia organization remained comparable to controls. Apart from that, Kl deficiency has been associated with increased soft tissue and vascular calcification in several organs [31]. Furthermore, in humans, calcification within the CP increases with age [32], suggesting that both aging and loss of Klotho may contribute to mineral deposition in this tissue. To investigate this potential relationship, we performed Von Kossa staining to assess whether Klotho deletion affects calcification in the CP. Our results indicated that Kl deletion differentially affects calcification in the two CP regions: it promoted early and more pronounced calcification in the FV-CP of young mice, while in the LV-CP, calcification increased mainly in aged mice. These region-specific differences may be explained by the distinct structural characteristics of the two CPs, as the FV-CP is more densely folded and layered compared to the more delicate LV-CP [9], potentially making it more resistant to structural disruption but more prone to retaining mineral deposits. Such anatomical variations likely contribute to the distinct responses of each CP region to aging and Kl deletion.

Furthermore, CP-specific Kl deletion had no impact on the aging-related phenotypic and behavioral changes observed in aged mice. Kl^ΔCP^ mice showed no significant differences in body weight, serum calcium or phosphate levels, brain/body weight ratio, or behavioral performance compared to controls. This indicates that CP-specific Kl loss does not cause systemic alterations or overt neurological deficits. Consistent with this, proteomic analysis revealed no single protein consistently altered across both CP regions in Kl^ΔCP^ mice, suggesting the presence of compensatory mechanisms that maintain CP homeostasis. A reason for that might also have been a large intra-group heterogeneity in some of the age groups. Such compensation may, at least in part, arise from circulating factors influenced by renal Klotho, as previous studies have shown that kidney-derived Klotho, although unable to cross the blood–brain barrier, can exert indirect effects on the brain through mediators such as platelet factor 4 (PF4) [33]. In addition, local compensatory responses within the CP itself, or in interconnected brain regions such as the hippocampus, may contribute to the activation of protective pathways that counterbalance the loss of Klotho in the CP.

Nevertheless, when aging was combined with Kl deletion, subtle molecular changes became apparent in the LV-CP, particularly in pathways related to extracellular matrix (ECM) remodeling, cytoskeletal organization, and protein homeostasis. The increased expression of integrins (Itga7, Itgb6, Tln1, Ptpra) and laminins (Lamc1, Lama1, Lama5) suggests enhanced cell–ECM interactions and activation of integrin-mediated signaling, which may promote ECM remodeling and create a microenvironment favorable for calcification. In parallel, the upregulation of cytoskeletal and junctional proteins (Pxn, Coro1c, Gas7, Fmn13, Spta1, Cdc42ep4, Cdc42se2, Phip, Anxa1) could influence cell shape and membrane organization in ways that facilitate mineral deposition. Although a direct causal link has not yet been demonstrated, previous studies indicate that some of these proteins—particularly Annexin A1, which binds calcium—may contribute to or promote mineralization processes [34], consistent with the increased calcification observed in the LV-CP of older Kl^ΔCP^ mice.

In both FV-CP and LV-CP, a decline in protein-folding and proteostasis-related pathways was observed, highlighting Kl’s important role in maintaining protein homeostasis during aging. The regional differences observed suggest that Kl deletion influences aging pathways in a subregion-specific manner, with the LV-CP appearing more vulnerable to subtle, age-dependent alterations.

In conclusion, while Kl deletion in the CP does not fully mimic natural aging, it partially activates similar molecular pathways, particularly those involved in ECM remodeling, cytoskeletal dynamics, and proteostasis.

Although this study provides new insights into the regional and molecular effects of Kl deletion in the CP, it is not yet complete. Additional experiments are needed to further strengthen and extend these findings. In particular, assessing behavioral performance at older ages would help clarify whether the cognitive effects of Kl deletion become more pronounced with aging. Moreover, the impact of CP-specific Kl deficiency may be more evident when combined with other pathological conditions, such as Alzheimer’s disease models or experimentally induced inflammation, which could better reflect the complex interactions occurring in neurodegenerative and age-related contexts. Furthermore, as the CP is the primary site of cerebrospinal fluid (CSF) production and secretion, analyzing the CSF could provide valuable insights into how the absence of Kl influences CSF composition and secretion dynamics. Such analysis may also reveal how Kl deficiency in the CP indirectly affects other brain regions through CSF-mediated alterations in the brain’s microenvironment.

## Materials and methods

### Generation of CP-Specific Klotho Knockout Mice

Klotho-flox mice (Kl^fl/fl^) were generated by Taconic Artemis (project FLI0005) by flanking exon 2 of the Kl gene with loxP sites. When Cre-recombinase is activated, exon 2 is excised, causing a frameshift in the Kl coding sequence, resulting in a truncated, nonfunctional protein. To generate CP-specific Kl knockout mice, we crossed Kl^fl/fl^ mice with Slco1c1-creERT2 mice that carry a tamoxifen-inducible Cre recombinase gene within the Slco1c1 gene locus [35]. Slco1c1 encodes for the organic anion transporter OATP1C1 that is highly expressed in capillary endothelial and CP epithelial cells in murine CNS [36]. As Kl is not present in CNS endothelial cells, cre-recombinase activation in the Kl^fl/fl^:Slco1c1-creERT2 animals is expected to inactivate Kl expression specifically in choroid plexus epithelial cells. Additionally, mice harboring a YFP reporter cassette in the ROSA26 locus were crossed in to monitor Cre-recombinase activity. For this study, we used three experimental groups with the genotypes: Kl^fl/fl^;Slco1c1-creER^+/+^;YFP^tg/tg^, as control#1 (Ctrl1); Kl^+/+^;Slco1c1-creER^tg/+^;YFP^tg/tg^, as control#2 (Ctrl2); and Kl^fl/fl^;Slco1c1-creER^tg/+^;YFP^tg/tg^, as the experimental knockout (Kl^ΔCP^) group. All experiments were conducted exclusively with female mice.

### Tamoxifen induction

TAM solution was prepared by dissolving 100 mg of TAM (Sigma, T5648) in 0.5 ml of 100% ethanol, followed by vigorous vertexing. Next, 9.5 ml of corn oil (Sigma, C8267) was added, and the mixture was heated at 55 °C for 1 hour with occasional shaking to ensure complete dissolution, resulting in a final concentration of 10 mg/ml. The solution was aliquoted and stored in light-protected tubes at −20 °C. Before injection, the solution was brought to room temperature and mixed. Each animal received 1 mg/100 µl of the solution intraperitoneally using a 1 ml syringe with a G26 or G27 needle. Injections were administered near the inner thigh to avoid organ damage. The injections were administered daily for 5 consecutive days on 4-5 week old mice, followed by a one-week pause, and then repeated for an additional 3 days.

### Genotyping

For genotyping, genomic DNA was extracted from tail biopsies by incubating them in 250 µl of Direct PCR (Tail) buffer (PeqLab, 31-102-T) with the addition of 3.75 µl of Proteinase K solution (VWR, 1245680500) overnight at 55°C in a thermomixer set to 800 rpm. After lysis, the PCR reaction was performed. Details of the primers and PCR conditions are provided in Supplementary Table 1 and 2.

### Histology and Immunostaining

Brain coronal sections, 20 µm thick, were prepared using a Leica CM3000 cryostat-microtome (Germany). After thawing, slides were air-dried under controlled conditions. Fixation was carried out using 4% Roti Histofix for 10 minutes with gentle agitation, followed by three 10-minute PBS washes at room temperature (RT). Permeabilization was achieved using 0.03% SDS in PBS for 3 minutes, followed by three more PBS washes. The sections were blocked with 1% bovine serum albumin (BSA) in PBS-T (0.2% Triton X-100) for 1 hour at RT. A primary antibody solution, prepared in PBS-T with 1% donkey serum, was applied (75 µl per section), and the slides were incubated overnight at 4°C. Following three 15-minute PBS washes, secondary antibody solution containing Hoechst 33258 (1:10,000), 1% blocking serum, and secondary antibody (1:1,000) was applied. After 2-3 hours of incubation at RT, slides were washed again and mounted using Fluoromount™ and glass coverslips. Images were captured using 20X or 63X objectives on an Axiovert 200 ApoTome microscope, controlled by ZEN 3.7 software. The Z-stack function was utilized to acquire images across multiple layers, which were then compiled using the tile function and maximum intensity projection.

### H&E and Von Kossa Staining

For H&E staining, cryosections were air-dried at room temperature for at least 20 min. Staining was performed using a Leica Stainer XL automated stainer with the following program: two initial washing steps with freshly prepared tap water (1 min each), hematoxylin solution (Gill II, staining solution; Morphisto, 1 min), a subsequent wash (3 min), and eosin solution (0.1% aqueous staining solution; Morphisto, 30 s). The sections were then rinsed three times with freshly prepared tap water (1 min each), dehydrated through graded ethanol solutions (95% and 100%, two changes each for 1 min), cleared in xylene (clearing reagent; Morphisto, two changes, 1 min each), and immediately coverslipped using the Leica CV5030 coverslipper with a xylene-based mounting medium (CV Mount, Leica). For Von Kossa staining, the Abcam kit (ab150687) was used. First, slides containing the brain sections were incubated in 5% silver nitrate solution for 20 minutes under ultraviolet light (250-280 nm) in a cell culture hood. After incubation, the slides were rinsed with distilled water, followed by a 2–3 minute incubation in a 5% sodium thiosulfate solution. Next, they were rinsed in running tap water for 2 minutes, followed by additional washes in distilled water. Slides were then incubated in Nuclear Fast Red solution for 5 minutes, washed again, and dehydrated in absolute alcohol. Finally, slides were cleared, mounted with Fluoromount™, and covered with glass coverslips. Images of the LV-CP and FV-CP were acquired using a Zeiss Axio Scan.Z1 slide scanner. Quantification of calcification was performed by first segmenting the respective CP regions using the Segment Anything AI-based server (https://segment-anything.com/demo) to accurately isolate the tissue of interest. The proportion of calcium deposits relative to the total tissue area was subsequently analyzed using a custom Python script executed via the Anaconda Prompt environment. Comparative analyses of calcification were conducted between Kl^ΔCP^ and control mice in both young (2–3 months old) and aged (20–21 months old) groups. The final data were analyzed using a two-sample *t*-test assuming unequal variances in Microsoft Excel (version 2403)

### Tissue Isolation and Serum Biochemistry Analysis

Mice were euthanized in a carbon dioxide (CO_2_) chamber with a controlled flow rate of 0.5 L/min, followed by aseptic dissection to collect various tissues, including blood, the lateral and fourth ventricle CP, hippocampus, tibialis anterior (TA) muscle, and kidney. Blood was collected via heart puncture, and serum was separated using Microvette® 500 Serum Gel (Sarstedt) by centrifuging for 5 minutes at 10,000g. The serum samples were then sent to the LABOKLIN Research Team (https://laboKlin.com/en/) for calcium and phosphate level analysis. As control, serum from 4-5 weeks Kl hypomorphic mice [1] was taken.

### Western blot Analysis

Protein extraction and quantification, SDS-PAGE and Western blot analysis were conducted as described previously [37]]. 15 µg of CP lysate was used for the Western blot analysis.

### Behavioral Tests

#### IntelliCage

The learning behavior and cognition of the mice were evaluated using the IntelliCage system, a standard polycarbonate cage with dimensions of 55 cm in width, 38 cm in length, and 21 cm in height. This specialized cage was equipped with four triangular operant test chambers, each measuring 15 cm × 15 cm × 21 cm, positioned at each corner. Additionally, round apertures on the walls of each chamber provided free access to water bottles. Prior to the experiment, each mouse was individually identified with subcutaneously injected radio-frequency identification (RFID) tags. These RFID tags allowed for automatic detection by a ring antenna at each chamber’s entrance, facilitating the recording of individual visits to each chamber. The mice were organized into groups of 10 and each group was placed in a separate IntelliCage for testing. The study was conducted over 17 days, comprising four phases: Free Adaptation, Nose-Poke Adaptation, Place Learning, and Retraining (Re-learning). During the Free Adaptation phase (7 days), the mice were allowed unlimited access to water, with the gates fully open. In the Nose-Poke Adaptation phase (3 days), access to water was contingent on nose-poking against a locked gate. The Place Learning phase (4 days) required the mice to access drinking water only in one of the four corners of the cage, necessitating nose-poking to open the gate and testing the mice’s ability to learn and remember the specific water source location. The Retraining phase (3 days) involved relocating the water source to a different corner, requiring the mice to adapt and learn the new location, with access once again nose-poke dependent. Throughout the 17 days, various metrics related to learning behavior were analyzed, with a primary focus on the Relearn Score. This score was calculated using the formula: Relearn Score = ln (Place Errors / Reversal Errors). A higher ratio indicated a faster relearning process. Negative values suggested fewer visits to unassigned corners during the Place Learning phase, reflecting effective learning behavior.

### Rotarod, Morris Water Maze, open field, hot plate, tail flick, and grip strength experiments

Spatial learning and memory were evaluated using the water maze task, as described in [21]. Motor performance was assessed using the accelerating rotarod test (TSE, Bad Homburg, Germany). Mice underwent a habituation session on day 1, followed by four days of trials in which the latency to drop from the rotating rod was measured. The rod had continuous rotation acceleration from 4 to 40 rpm over a period of 5 minutes. Pain sensitivity was measured using the hot plate test (Ugo Basile, Italy) and tail flick as described in [38]. The open field test was performed as described in [39].

Motor function and muscular strength were assessed using a grip strength test (TSE Systems). Two types of grids were used to measure forelimb and all-limb strength. The peak force exerted before release was recorded, with each mouse tested four times.

### LC-MS Proteomics Analysis

The CP and hippocampus tissues were processed for LC-MS proteomics analysis. Tissues were resuspended in PBS and a lysis buffer containing 5% SDS, 100 mM HEPES, and 50 mM DTT. Samples were sonicated using a Bioruptor Plus (Diagenode, Belgium) for 10 cycles (30 sec ON/60 sec OFF) at a high setting and 20°C. Following sonication, the samples were boiled at 95°C for 7 minutes. Subsequent steps followed a previously described protocol [37].

### Statistical Analyses for behavioral tests

To assess the equality of variance, we applied Levene’s test before the analysis of variance (ANOVA) [40] to verify equality of variances across groups. In some cases, Q-Q plots were generated to compare data distribution against a theoretical normal distribution. Repeated Measures ANOVA (RM-ANOVA) was used to examine mean differences across time points and genotypes. A p-value of less than 0.05 was chosen as a significance threshold. Additionally, Cohen’s d effect size was used in post hoc analyses to quantify the magnitude of differences between experimental conditions, providing a comprehensive interpretation of both statistical significance and practical relevance.

## Acknowledgments

We gratefully acknowledge the support of the animal, imaging, histology, and proteomics core facilities at the Leibniz Institute on Aging – Fritz Lipmann Institute (FLI). We thank Jana Hamann and Daniela Reichenbach for excellent technical assistance. We also thank Jihen Amara for her valuable contribution to the development of Python scripts used in the data analysis.

## Author contributions

Conceptualization, C.K.; validation, Z.F.-K, A.G., C.V., R.B., H.H., M.S.; formal analysis, Z.F.-K., T.P., S.H, R.B. and A.G.; investigation, Z.F.-K.; writing—original draft preparation, Z.F.-K.; writing—review and editing, Z.F.-K. and C.K. T.P., and C.V.; visualization, Z.F.-K. T.P., and A.G.; supervision, C.K.; project administration, C.K.; funding acquisition, C.K. All authors have read and agreed to the published version of the manuscript.

## Institutional Review Board Statement

Breeding, organ removal and animal experiments were performed under the general license of the institute (TG/3-J-0002858/V-98/23) and the animal licence FLI-20-009, both granted by the local authority, the Thüringer Landesamt für Verbraucherschutz.

