## Supplemental material for "Distinct Effects of Aging and Klotho Deletion on the Choroid Plexus"

**Supplementary Table 1. List of primers used for genotyping of newly born pups**

| Locus | Primer name | Primer sequence | Expected product size | PCR Program |
|---|---|---|---|---|
| **Slco1c1-Cre** | Alfi cre<br>Alfi cre rev<br>B2-1<br>B2-2 | 5’-AGA TGC CAG GAC ATC AGG AAC CTG-3’<br>5’-ATC AGC CAC ACC AGA CAC AGA GAT C-3’<br>5’-CAC CGG AGA ATG GGA AGC CGA A-3’<br>5’-TCC ACA CAG ATG GAG CGT CCA G-3’ | +/+: 300bp<br>tg/+: 250bp + 300bp<br>tg/tg: 250bp | 1. 94 °C - 3'<br>2. 94 °C - 30''<br>3. 60 °C - 30''<br>4. 72 °C - 1'<br>5. 72 °C - 10'<br>6. 10 °C - hold<br>Step 2.-4. 35x |
| **Klotho-flox** | Kl 33_fwd<br>Kl 34_rev | 5’-TAT GTA GAG ?GG ?TT TAC AAG G-3’<br>5’-GTC TTC CCT GTG AAT CAG CC-3’ | +/+: 195 bp<br>fl/+: 195+305bp<br>fl/fl: 305 bp | See above |
| **YFP** | oIMR4982 Mutant<br>oIMR8545 common<br>oIMR8546 wt | 5’-AAG ACC GCG AAG AGT TTG TC-3’<br>5’-AAA GTC GCT CTG AGT TGT TAT-3’<br>5’-GGA GCG GGA GAA ATG GAT ATG-3’ | +/+: 600bp<br>tg/tg: 320 bp<br>tg/+: 320+600 bp | See above |

**Supplementary Table 2. Reaction mix for genotyping of newly born pups**

| Component | for 20 µl reaction | Supplier |
|---|---|---|
| **5x GoTaq reaction buffer** | 4 µl | Promega, M7911 |
| **10x dNTPs** | 2 µl | New England Biolabs, N0447S |
| **GoTaq polymerase** | 0.1 µl | Promega, M7848 |
| **Each primer** | 0.5 µl | Eurofins Genomics |
| **Genomic DNA** | 0.5 µl | |
| **ddH2O** | Up to 20 µl | |

**Supplementary Table 3. Differentially expressed proteins across different ages in FV-CP (Combined data from comparing Ctrl1 to $Kl^{\Delta CP}$and Ctrl2 to $Kl^{\Delta CP}$)**

| Tissue/Age | number of proteins | Gene symbol |
|---|---|---|
| FV-CP/M14-15, M2-3, M5-6 | 1 | Abca5 |
| FV-CP/M2-3, M5-6 | 2 | Ndufb9, Mgrn1 |
| FV-CP/M14-15, M5-6 | 1 | TCAIM |

| | | | | |
|---|---|---|---|---|
| FV-CP/M2-3 | 35 | *Mp68* | *D2hgdh* | *Dynlt1* |
| | | *Lifr* | *Mepce* | *Vtn* |
| | | *Banf1* | *Mpdu1* | *Rsrc2* |
| | | *Abcg2* | *Fuom* | *Mog* |
| | | *Retsat* | *Sugp1* | *Trappc4* |
| | | *Jph1* | *Ppl* | *Eps15* |
| | | *Serpina3k* | *Sfr1* | *Adgrl1* |
| | | *Slc7a1* | *Tmem27* | *Ceacam2* |
| | | *Casp6* | *Sphk1* | *Tmem186* |
| | | *Sema7a* | *C1qc* | *Atg12* |
| | | *Dscr3* | *Arl8a* | *Hrnr* |
| | | *Hspb11* | *H3f3c; H3f3a* | |
| FV-CP/M5-6 | 53 | *Coch* | *Gstp1* | *Hspa1b; Hspa1a* |
| | | *Fabp5* | *Gstm5* | *Cox6a1* |
| | | *Atp5f1* | *Pcp4* | *Atp8a2* |
| | | *Mettl13* | *Rab33b* | *Ifih1* |
| | | *Scpep1* | *Cfd* | *Abi3* |
| | | *Dpp4* | *Wiz* | *Sat2* |
| | | *ALB* | *Znf593* | *Rpl38* |
| | | *Idh3g* | *Sdhaf2* | *Atp2a1* |
| | | *Ptgds* | *Cdkal1* | *Atp1b3* |
| | | *Lsm3* | *Prr14* | *Csnk2a1* |
| | | *Nptn* | *Ndufs8* | *Alg10b* |
| | | *Ypel5* | *Mcts1* | *Mrpl49* |
| | | *Wdr77* | *Crabp2* | *Tcp11l1* |
| | | *Fgfr1* | *Col6a1* | *Aatf* |
| | | *Fam136a* | *Hspa14* | *Thap4* |
| | | *Adam9* | *Trmt6* | *Mrps30* |
| | | *Taco1* | *Col6a2* | *Nt5c3b* |
| | | *Tmem263* | *Tmem209* | |
| FV-CP/M14-15 | 10 | *Slc5a6* | *Aldh3a1* | |
| | | *Nudt6* | *Nfatc2* | |
| | | *Atpaf1* | *Znf207* | |
| | | *HVM51* | *Idua* | |
| | | *Emc6* | *Tubb1* | |
| FV-CP/M20-21 | 15 | *Naaa* | *Apob* | |
| | | *Baiap2l1* | *Lamp2* | |
| | | *Gm11992* | *Ubap1* | |
| | | *Nrm* | *Cfl2* | |
| | | *Ifit3* | *Smad2* | |
| | | *Mrpl52* | *Efemp2* | |
| | | *Rad23a* | *Zmat2* | |
| | | *Mtnd5* | | |

**Supplementary Table 4. Differentially expressed proteins across different ages in LV-CP (Combined data from comparing Ctrl1 to Kl$^{\Delta CP}$and Ctrl2 to Kl$^{\Delta CP}$)**

| Tissue/Age | number of proteins | Gene symbol |
|---|---|---|
| LV-CP/M2-3, M5-6 | 1 | *Sult1c2* |

| | | | |
|---|---|---|---|
| LV-CP/M2-3 | 40 | *Fbln5*<br>*Utp15*<br>*Lsm4*<br>*Cadm1*<br>*Osbpl3*<br>*Phf24*<br>*Thy1*<br>*Cnp*<br>*Fam241b*<br>*Gna11*<br>*Ptgr1*<br>*Aldh3a2*<br>*Igh-3*<br>*Mag*<br>*Thap4*<br>*Hat1*<br>*Dnajc5*<br>*Plcb1*<br>*Abca5*<br>*Nefl* | *Dnm2*<br>*Ndrg1*<br>*Cndp1*<br>*Alcam*<br>*Rbmx*<br>*Smap1*<br>*Cyp4v2*<br>*Hmgcl*<br>*Slc7a10*<br>*St3gal6*<br>*Srr*<br>*H2afy2*<br>*Ptprd*<br>*Rhbdf2*<br>*Ctnna2*<br>*C1qc*<br>*Fetub*<br>*Tmem70*<br>*Cpsf3l*<br>*Tbl1x* |
| LV-CP/M5-6 | 17 | *Fuz*<br>*Ndel1*<br>*Sap30bp*<br>*Zfpl1*<br>*Ap3b2*<br>*Cln8*<br>*Gja1*<br>*Fabp7*<br>*Syt1* | *Nos1ap*<br>*Tspo*<br>*Camk2b*<br>*Il6st*<br>*Syn2*<br>*Ngly1*<br>*Tcp11l1*<br>*Phlda3* |
| LV-CP/M14-15 | 34 | *Rpl35*<br>*Nme1*<br>*Sdpr*<br>*Csrp1*<br>*Tubb6*<br>*Hnrnpm*<br>*Fn3krp*<br>*Hist1h1d*<br>*Uqcrb*<br>*Tbc1d24*<br>*Alpl*<br>*Hn1l*<br>*Dysf*<br>*Chp1*<br>*Rwdd1*<br>*Mrc1*<br>*Cd63* | *Ddr1*<br>*Capza1*<br>*Steap3*<br>*Pcp4*<br>*Lamc3*<br>*Mavs*<br>*Fam103a1*<br>*Hdgf*<br>*Rps27l*<br>*Tubb2b*<br>*Arf5*<br>*Eef1a1*<br>*Arpc5*<br>*Ahcyl2*<br>*Rbmxl1*<br>*Pacrg*<br>*Rps10* |
| LV-CP/M20-21 | 5 | *Col3a1*<br>*Fbn1*<br>*Arpp19* | *Col1a2*<br>*Pvalb* |

**Supplementary Table 5. KEGG pathway analysis of the proteins downregulated (in all ages) in Kl$^{\Delta CP}$ mice versus controls in the HC**

| Term | Adjusted P-value | Protein symbol |
|---|---|---|
| **Parkinson disease** | 6.24E-04 | Gnal; Ndufa4; Camk2a; Calm1; Tubb4a |
| **Long-term potentiation** | 0.002421 | Camk2a; Araf; Calm1 |
| **Glioma** | 0.002421 | Camk2a; Araf; Calm1 |
| **Pathways of neurodegeneration** | 0.003417 | Ndufa4; Camk2a; Araf; Calm1; Tubb4a |
| **Dopaminergic synapse** | 0.007738 | Gnal; Camk2a; Calm1 |
| **Alzheimer disease** | 0.010469 | Ndufa4; Araf; Calm1; Tubb4a |

**Supplementary Table 6. KEGG pathway analysis of the proteins upregulated (in all ages) in Kl$^{\Delta CP}$ mice versus controls in the HC**

| Term | Adjusted P-value | Protein symbol |
|---|---|---|
| **ECM-receptor interaction** | 1.80E-08 | Col1a1; LAMA5; Lamb2; Col4a1; Lamc1; Hspg2 |
| **Amoebiasis** | 2.21E-08 | Col1a1; Lama5; Gnal; Lamb2; Col4a1; Lamc1 |
| **Focal adhesion** | 2.69E-05 | Col1a1; Lama5; Lamb2; Col4a1; Lamc1 |

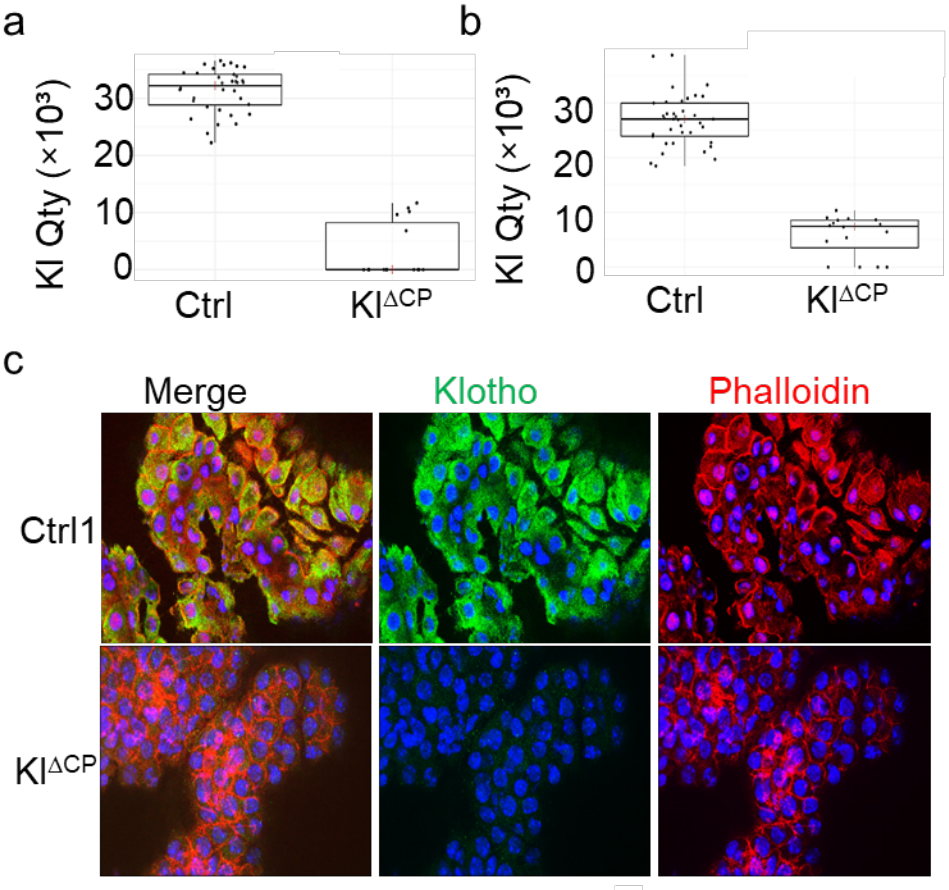

**Supplementary Figure 1. Proteomics analysis and Immunofluorescent staining and confirmed the deletion of Kl in LV-CP of Kl$^{\Delta CP}$ mice.** (a, b) Validation of Kl knockout in the CP of Kl$^{\Delta CP}$ mice using LC/MS proteomics of the FV-CP (A, n=36 controls, n=15 Kl$^{\Delta CP}$) and the LV-CP (B, n=37 controls, n=16 Kl$^{\Delta CP}$). (c) Brain sections from 2-3 months old Kl$^{\Delta CP}$ and control 1 mice were immunostained for Klotho and Phalloidin, and Z-stack images were acquired using the tile scan function on a Zeiss ApoTome microscope. Maximum intensity projections are shown. Nuclei were counterstained with Hoechst (blue). Scale bar = 20 µm.

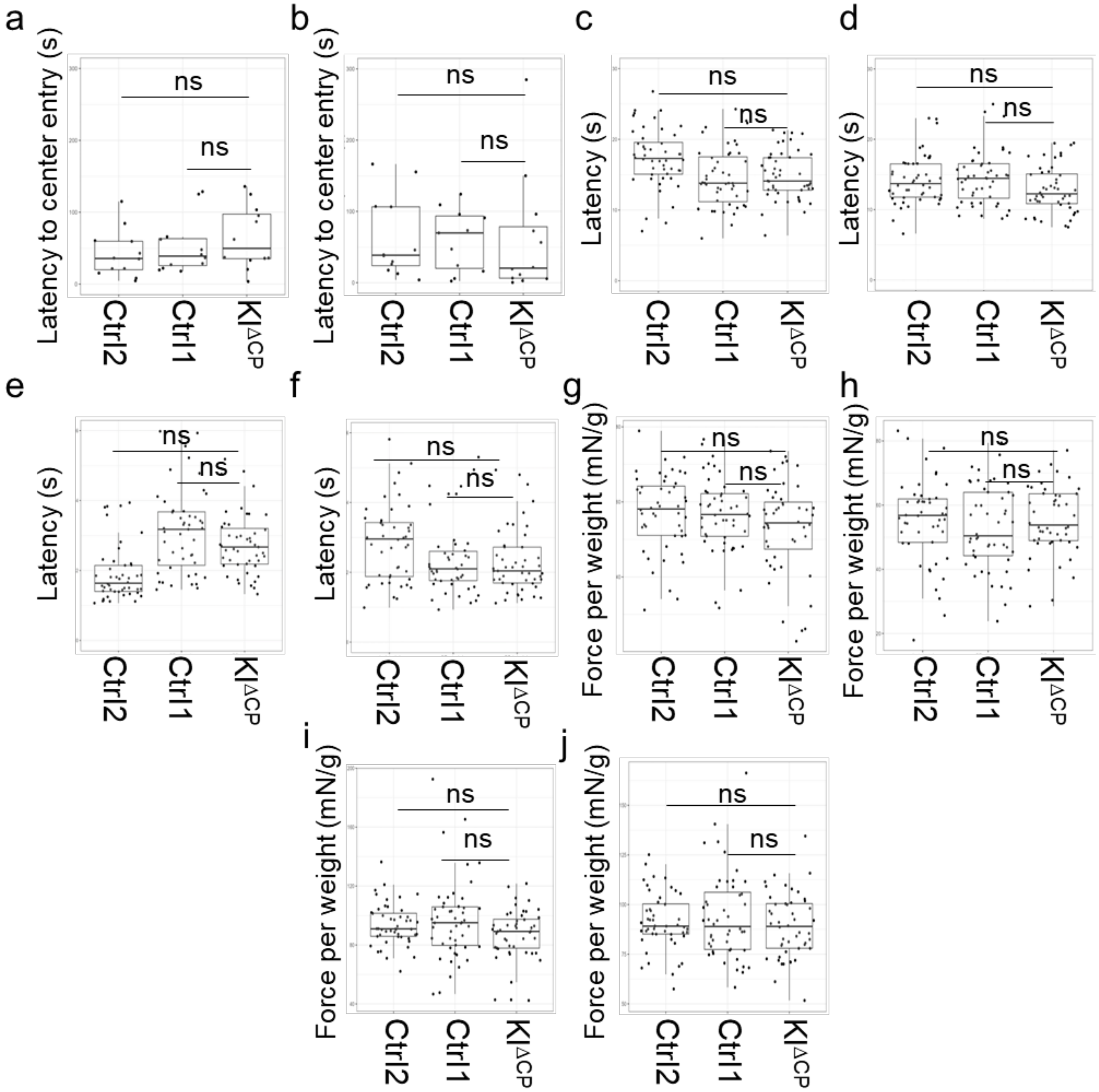

**Supplementary Figure 2. Behavioral analysis of 2-5 months and 8-12 months $KI^{\Delta CP}$ mice compared to control groups showed no significant difference between them.** (a, b) Open field test to measure exploration behavior (latency to center entry) between $KI^{\Delta CP}$ and control mice. The test involved placing the animal in a white plastic cube. Movement was tracked using a video system that was installed above the area. The latency to enter the center was assessed in (a) 2-5 months and (b) 8-12 months mice. (c, d) Hot plate test. Each mouse was placed on a 52°C preheated plate and the latency time until the occurrence of the first contact avoidance reaction was measured. Measurements were taken for (c) 2-5 months and (d) 8-12 months. (e, f) Tail flick response. In this experiment the mouse's tail of 2-5 months (e) and 8-12 months (f) was immersed in hot water at 45.0±0.2 °C and the reflexive tail movement time was recorded. (g-j) Grip strength analysis measuring the peak force exerted by front paws or all four paws on a wire grid. Data represents (g) 2-paw strength in 2-5 months, (h) 2-paw strength in 8-12 months, (i) 4-paw strength in 2-5 months, and (j) 4-paw strength in 8-12 months. Statistical evaluation included RM-ANOVA and Cohen's d-effect size, with error bars representing standard deviation and each dot symbolizing a single mouse. Significance was set at p-value > 0.05 (ns).

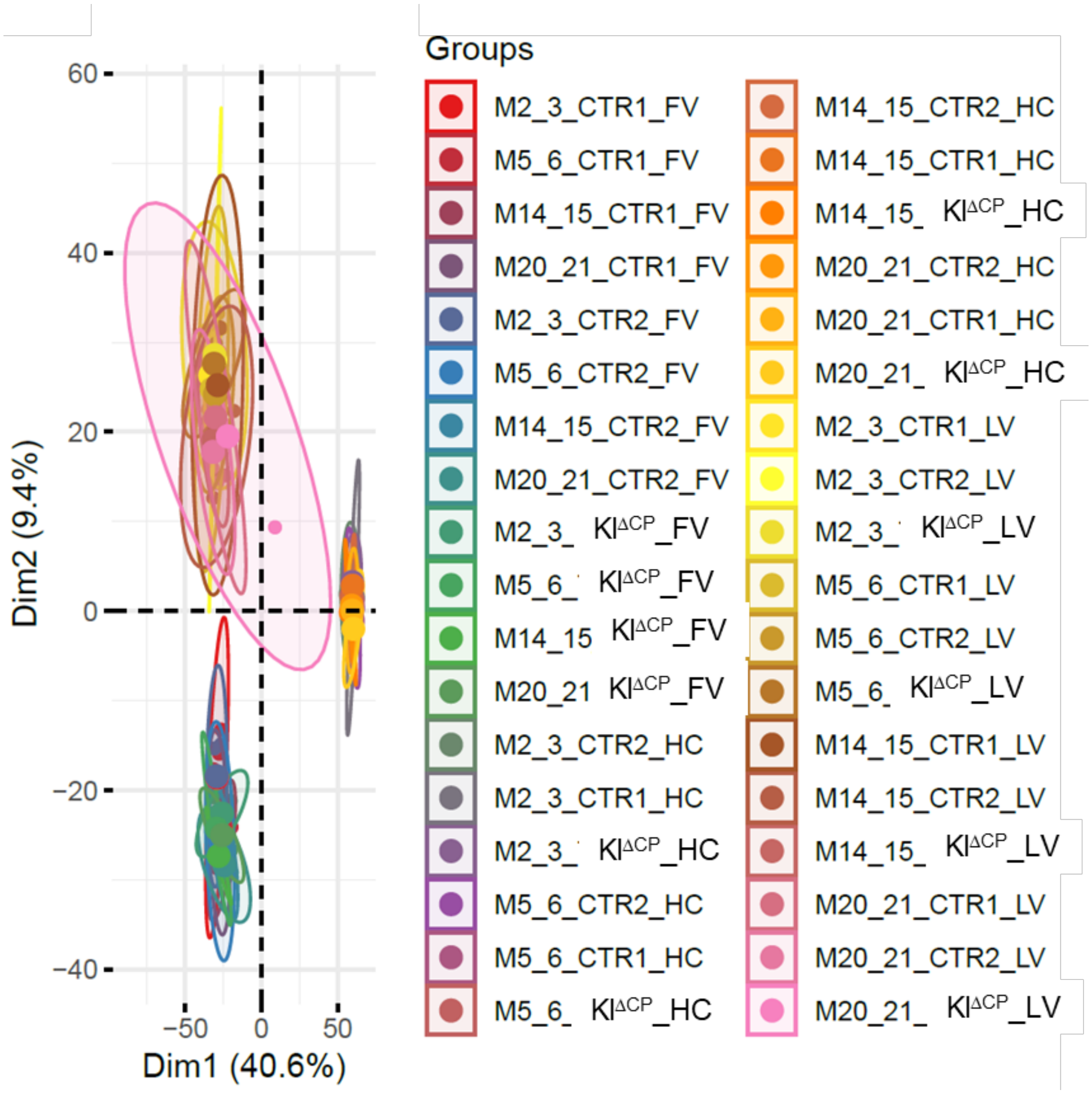

**Supplementary Figure 3.** Principal component analysis (PCA) illustratingthe distribution of all samples. The results show clear clustering of FV-CP, LV-CP, and HC samples within their respective groups, indicating reproducible replicate data and distinct proteomic profiles for FV-CP, LV-CP, and HC tissues (M: Month, CTR: Control, FV-CP: Fourth ventricle Choroid plexus, LV-CP: Lateral ventricle Choroid plexus, HC: Hippocampus).

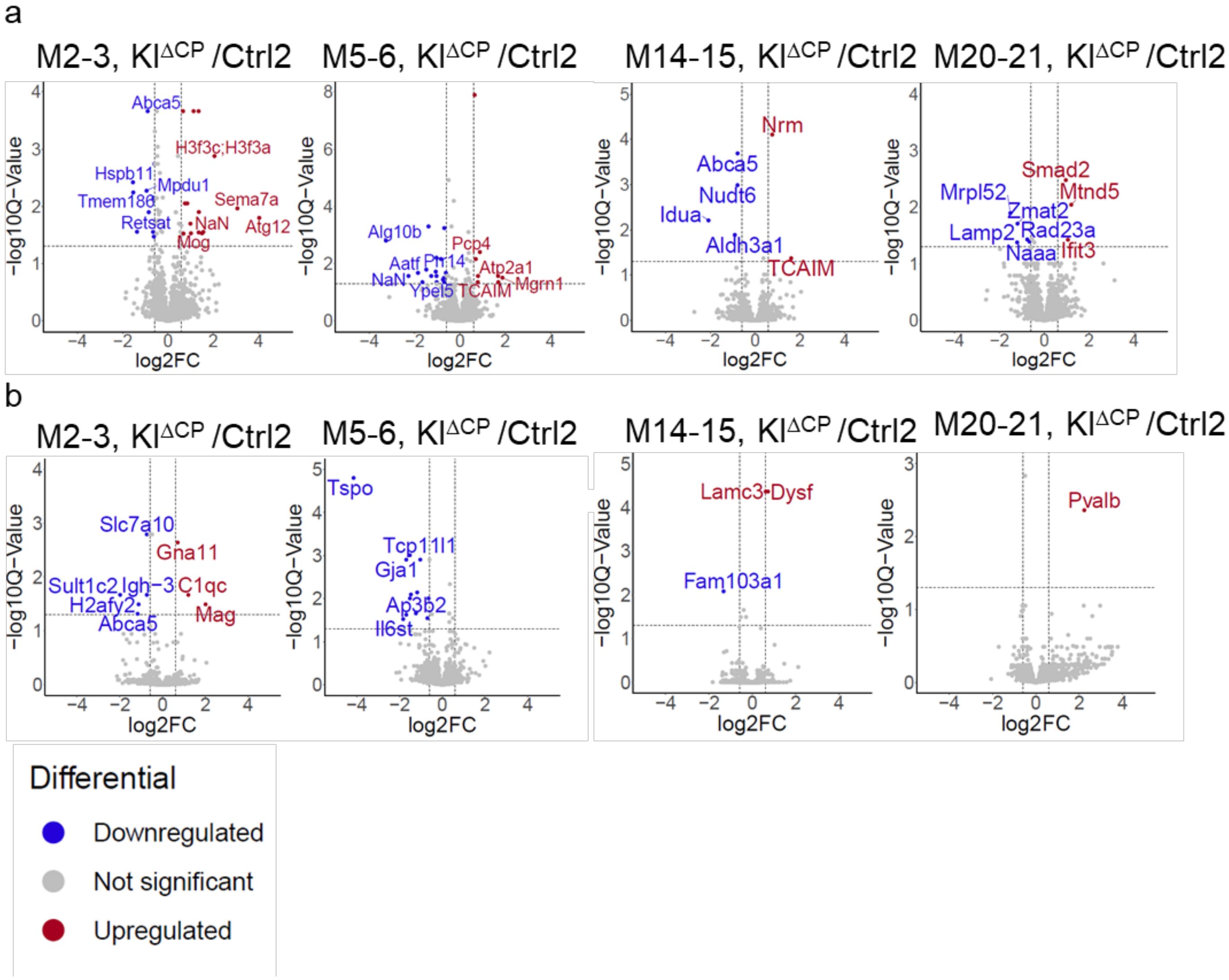

**Supplementary Figure 4. Volcano plots depicting differential expression of proteins in the FV-CP (a) and LV-CP (b) in KI$^{\Delta CP}$ mice compared to Ctrl2 at various ages (2-3, 5-6, 14-15, and 20-21 months).** Shown are protein symbols, the written names are the top 5 upregulated and downregulated proteins. Blue, significantly downregulated in KI$^{\Delta CP}$ compared to Ctrl2, red, upregulated in KI$^{\Delta CP}$ compared to Ctrl2.

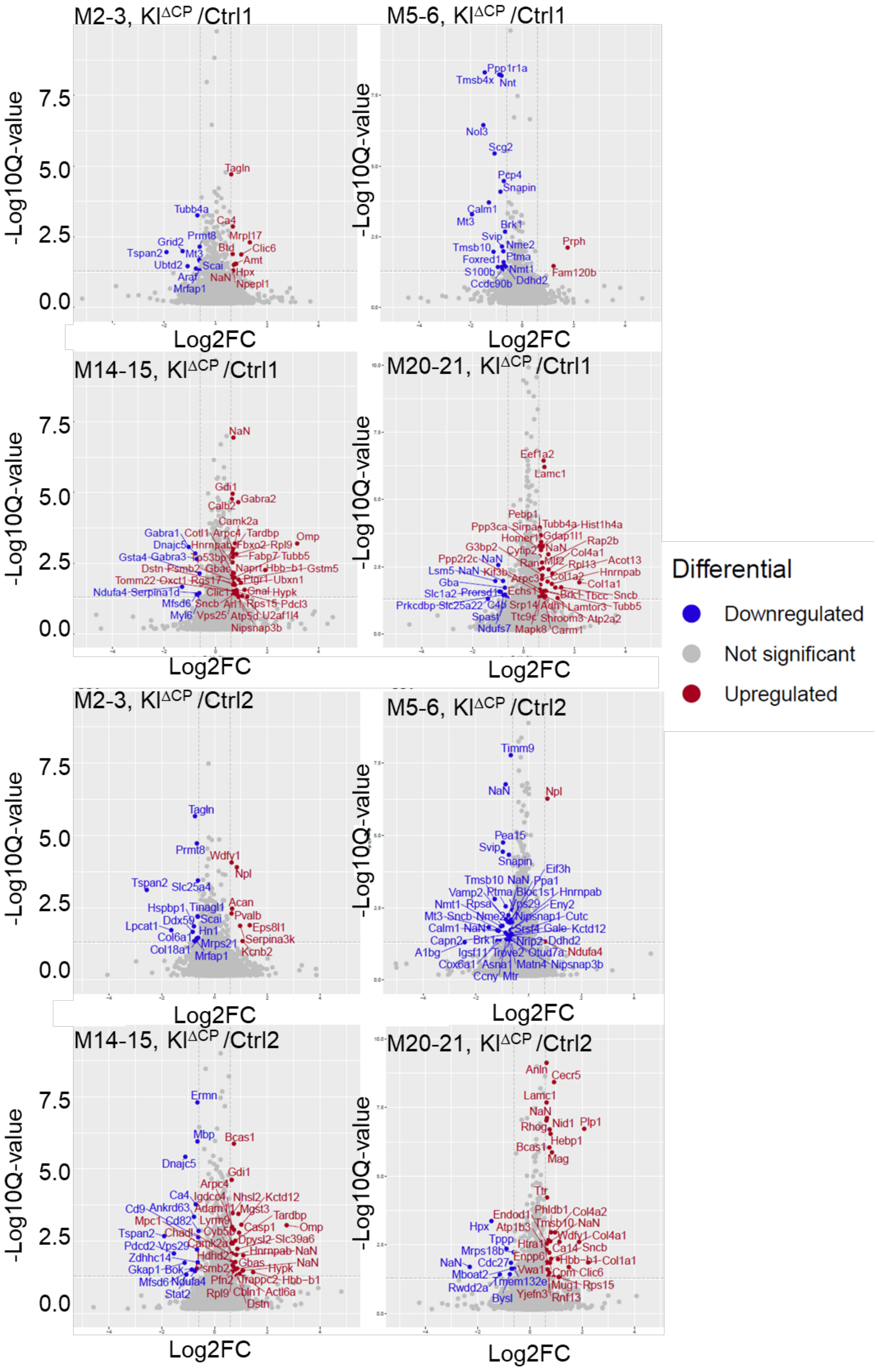

M2-3, KI$^{\Delta CP}$ /Ctrl1
M5-6, KI$^{\Delta CP}$ /Ctrl1
M14-15, KI$^{\Delta CP}$ /Ctrl1
M20-21, KI$^{\Delta CP}$ /Ctrl1
M2-3, KI$^{\Delta CP}$ /Ctrl2
M5-6, KI$^{\Delta CP}$ /Ctrl2
M14-15, KI$^{\Delta CP}$ /Ctrl2
M20-21, KI$^{\Delta CP}$ /Ctrl2
-Log10Q-value
Log2FC
7.5
5.0
2.5
0.0
Differential
Downregulated
Not significant
Upregulated
Tagln
Tubb4a
Ca4
Mrpl17
Prmt8
Grid2
Clic6
Tspan2
Mt3
Btd
Amt
Ubtd2
Scai
Hpx
Araf
NaN
Npepl1
Mrfap1
Ppp1r1a
Tmsb4x
Nnt
Nol3
Scg2
Pcp4
Snapin
Calm1
Brk1
Svip
Nme2
Prph
Tmsb10
Ptma
Foxred1
Nmt1
Fam120b
S100b
Ddhd2
Cdc90b
Gdi1
Gabra2
Calb2
Camk2a
Omp
Eef1a2
Lamc1
Pebp1
Timm9
Npl
Pea15
Snapin
Eif3h
Wdfy1
Slc25a4
Acan
Pvalb
Eps8l1
Serpina3k
Kcnb2
Mrfap1
Ermn
Mbp
Bcas1
Dnajc5
Anln
Cecr5
Nid1
Plp1
Hebp1
Mag
Ttr

**Supplementary Figure 5. Volcano plots depicting differential expression of proteins in the hippocampus in KI$^{\Delta CP}$ mice compared to Ctrl1 and Crl2 at various ages (2-3, 5-6, 14-15, and 20-21 months).** Shown are protein symbols, the written names are all the upregulated and downregulated proteins. Blue, significantly downregulated in KI$^{\Delta CP}$ compared to Ctrl, red, upregulated in KI$^{\Delta CP}$ compared to Ctrl.

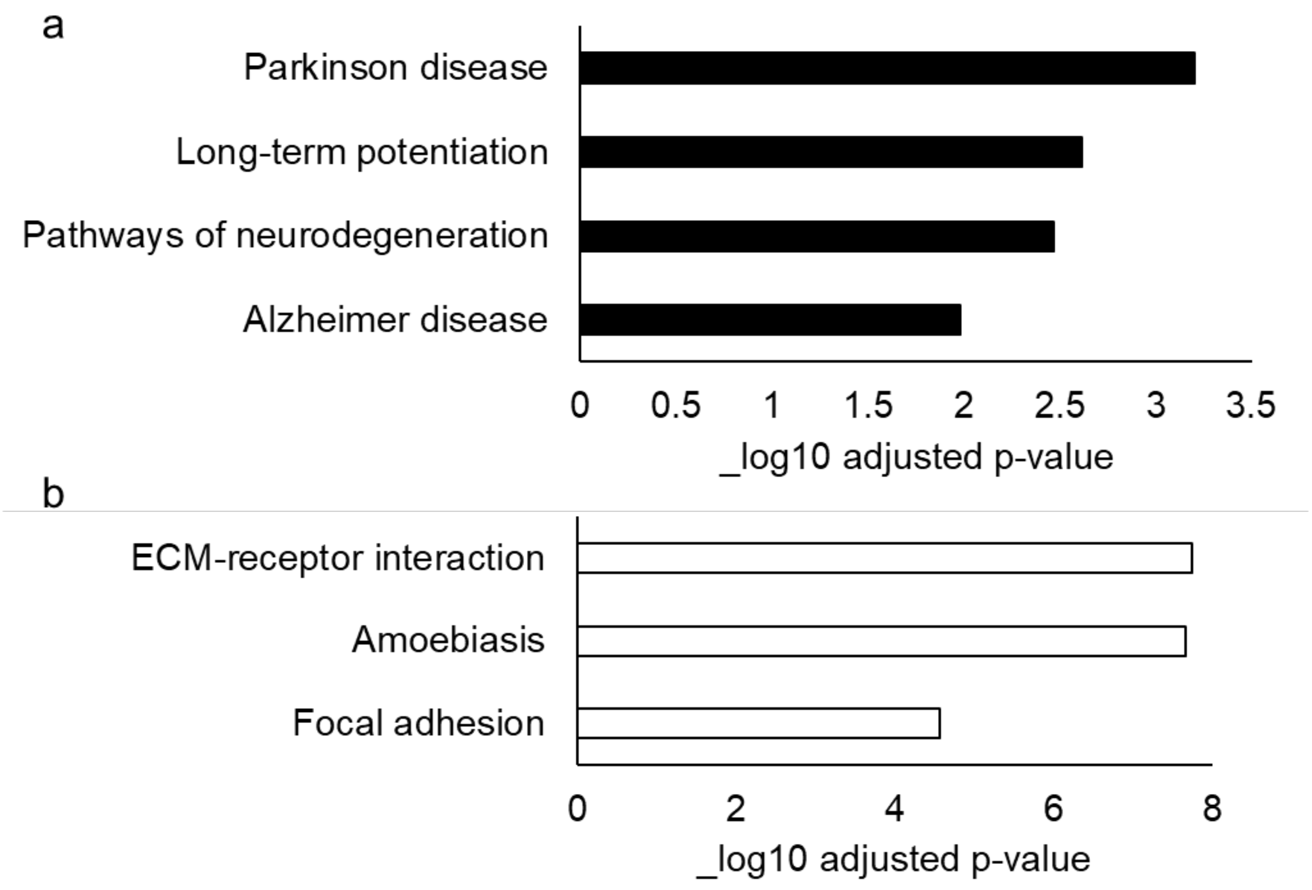

**Supplementary Figure 6.** KEGG pathway analysis of the proteins downregulated (a) and upregulated (b) (in all ages) in CP-KL KI$^{\Delta CP}$ mice versus controls in the HC.
